# Mural cell SRF controls pericyte migration, vessel patterning and blood flow

**DOI:** 10.1101/2021.11.26.470022

**Authors:** Michael Martin Orlich, Rodrigo Diéguez-Hurtado, Regine Muehlfriedel, Vithiyanjali Sothilingam, Hartwig Wolburg, Cansu Ebru Oender, Pascal Woelffing, Christer Betsholtz, Konstantin Gaengel, Mathias Seeliger, Ralf H. Adams, Alfred Nordheim

## Abstract

**Rationale:** Pericytes (PCs) and vascular smooth muscle cells (vSMCs), collectively known as mural cells (MCs), are recruited through PDGFB-PDGFRB signaling. MCs are essential for vascular integrity, and their loss has been associated with numerous diseases. Most of this knowledge is based on studies in which MCs are insufficiently recruited or fully absent upon inducible ablation. In contrast, little is known about the physiological consequences that result from impairment of specific MC functions.

**Objective:** Here, we characterize the role of the transcription factor serum response factor (SRF) in MCs and study its function in developmental and pathological contexts.

**Methods and Results:** We generated a mouse model of MC-specific inducible *Srf* gene deletion and studied its consequences during retinal angiogenesis. By postnatal day (P)6, PCs lacking SRF were morphologically abnormal and failed to properly co-migrate with angiogenic sprouts. As a consequence, PC-deficient vessels at the retinal sprouting front became dilated and leaky. By P12, also the vSMCs had lost SRF, which coincided with the formation of pathological arteriovenous (AV) shunts. Mechanistically, we show that PDGFB-dependent SRF activation is mediated via MRTF co-factors. We further show that MRTF-SRF signaling promotes pathological PC activation during ischemic retinopathy. RNA-sequencing, immunohistology, *in vivo* live imaging and *in vitro* experiments demonstrated that SRF regulates expression of contractile SMC proteins essential to maintain the vascular tone.

**Conclusions:** SRF is crucial for distinct functions in PCs and vSMCs. SRF directs PC migration downstream of PDGFRB signaling and mediates pathological PC activation during ischemic retinopathy. In vSMCs, SRF is essential for expression of the contractile machinery, and its deletion triggers formation of AV shunts. These essential roles in physiological and pathological contexts provide a rational for novel therapeutic approaches through targeting SRF activity in MCs.

## 1. Introduction

Blood vessels are composed of endothelial cells (ECs) lining the vascular lumen and surrounding mural cells (MCs) ^1^. MCs include pericytes (PCs), which cover capillaries, the smallest diameter blood vessels, and vascular smooth muscle cells (vSMCs)^2^, which cover arteries, arterioles, venules and veins, the larger caliber vessels ^3^.

During angiogenesis, new blood vessels develop from pre-existing ones by vascular sprouting. This process involves endothelial tip cells, specialized ECs that temporarily adopt a motile phenotype and extend numerous filopodia. Tip cells spearhead the sprouting vessels and secrete PDGFB, thereby attracting PCs that express the corresponding PDGFRB, to co-migrate along the nascent vessel sprout. PC recruitment is generally considered to aid vessel stabilization, and numerous studies have demonstrated the importance of PCs for vascular function ^4^. In the brain, PCs have been shown to express high levels of transporters and are considered to play a crucial role in maintaining homeostasis at the neurovascular unit (NVU)^3,5^. In accordance, failure to recruit PCs results in impaired formation of the blood-brain and the blood-retina barrier (BBB/BRB)^3^. Loss of PC coverage has been identified as an early event in diabetic retinopathy, which is associated with breakdown of the BRB ^6^. Most of our knowledge about PC function comes from studies in which PC recruitment has been compromised, or in which PCs have been ablated. Only few studies have investigate the physiological consequences that arise if specific aspects of PC biology or functions are inactivated ^7–10^. PC research is further complicated by the fact that identification and characterization of PCs is challenging due to the lack of unambiguous markers. Commonly used markers to identify PCs, such as PDGFRB, neural/glial antigen 2 (NG2), desmin or CD13, are also expressed by other cell types and therefore accurate identification of PCs requires, in addition, attention to their localization and morphology^11^.

In contrast to PCs, vSMCs are highly contractile and express a specific set of smooth muscle genes (SMG). Through a basal level of constriction, vSMCs create the vascular tone, which is essential to direct blood flow into capillaries ^5^. Vascular tone differs between organs and is dependent on the balance of competing vasoconstrictors and vasodilators ^12^. Dysregulation of vSMC contractility is associated with hypertension, aortic stiffness and chronic venous disease ^13–15^.

Serum response factor (SRF) is a conserved, ubiquitously expressed transcription factor which belongs to the MADS box protein family, and is known to regulate motile functions in a variety of cell types ^16–18^. SRF is activated either by Rho/actin or Ras/MAPK signaling ^17,18^. Those pathways involve different cofactor proteins, either myocardin related transcription factor (MRTF) or ternary complex factors (TCFs) and result in the expression of distinct sets of target genes. SRF has been reported to drive, amongst others, the expression of SMGs alpha-smooth muscle actin (*Acta2,* αSMA) and transgelin (*Togln,* SM22α) in visceral SMCs ^16,19^, but its role in pericytes and SMCs of the vasculature has not been thoroughly investigated. In order to address this question, we used *Pdgfrb-CreER^T2^* mice, which allow to delete SRF in PCs and SMCs of the vasculature ^7,20,21^ and, importantly, prevents the lethality associated with SRF deletion in visceral SMCs ^19,22^.

We demonstrate that SRF is crucial for distinct functional roles in PCs and vSMCs, respectively. In PCs, SRF is essential for cell migration downstream of PDGFRB signaling and mediates the pathologic activation of PCs during ischemic retinopathy. In vSMCs, SRF is essential for the expression of SMC genes, and its deletion triggers the formation of arteriovenous malformations (AVMs).

## 2. Methods

A detailed method section is provided by the supplementary section.

## 3. Results

### Conditional MC specific deletion of *Srf*

To address the role of SRF in MCs *in vivo,* we crossed floxed *Srf^flex1/flex1^* (meaning floxed **exon 1,** also referred to as *Srf^flox/flox^* mice) mice ^20^ with *Pdgfrb-CreER^T2^* mice, which have been shown to efficiently target MCs ^7,21^. We induced MC-specific deletion of *Srf* (hereafter referred to as *Srf^iMCKO^*) in newborn mice via tamoxifen administration at postnatal days (P) 1-3 and focused our analysis on the retinal vasculature. We choose three distinct time points (P6, P12 and after 4-8 weeks) for analysis in order to investigate the role of SRF during vascular sprouting, remodeling and maturity of the retinal vasculature (**Figure *1*, A-B**). To monitor recombination specificity and efficiency, we further introduced the *Rosa26^mTmG^* reporter^23^ to the *Pdgfrb-CreER^T2^::Srf^flox/flox^* mouse line. The *Rosa26^mTmG^* reporter expresses membrane-targeted GFP in cells, in which Cre mediated recombination has been occurred and revealed that, at P6, around 80% of mural cells had recombined **(Online Figure I, A-C).**

**Figure 1.**
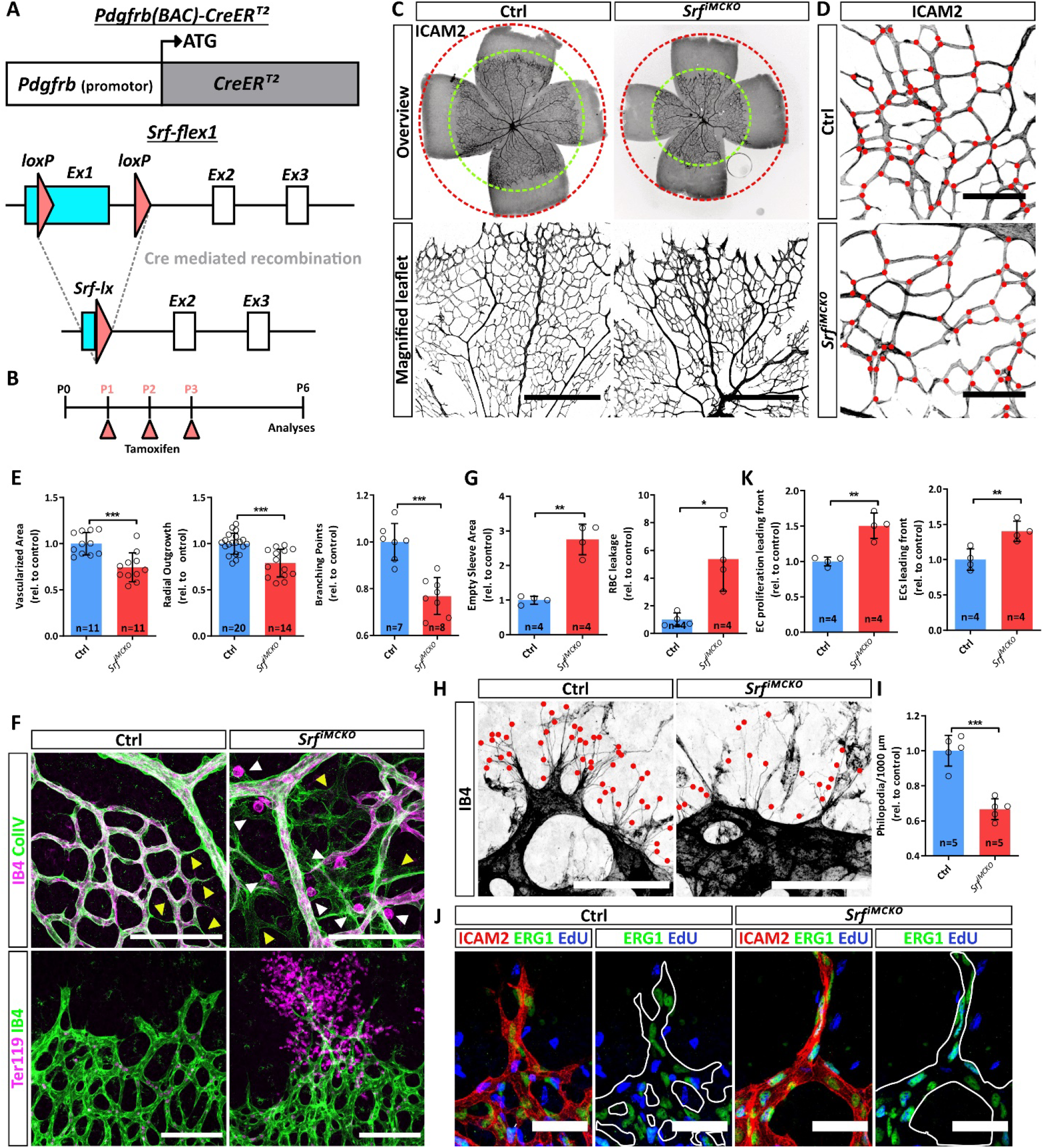
SRF-deficient Pericytes (PCs) impair early vascular morphogenesis of the retina. **(A)** Schematic representation of the *Pdgfrb(BAC)-CreER*^12^ transgene and Cre-mediated recombination of the *Srf-flex1* allele. (B) Tamoxifen administration regime and time point of analyses. (C) Epifluorescence overview images (upper panel) and confocal images (lower panel) of whole mount retinas stained for ICAM2 to visualize vascularization and vascular outgrowth of the primary vascular plexus. Scale bar, 250 μm. **(D)** Comparison of vascular branch points in control and *Srf^iMCKO^* retinas. Red dots indicate branch points. Scale bar, 100 μm **(E, G, K)** Quantifications of morphometric parameters, pruning, leakage and endothelial cell (EC) proliferation. **(F)** Confocal images of vascular pruning events visualized by empty Collagen IV sleeves (upper panel; yellow arrowheads) and vascular leakage, visualized as IB4 positive immune-cells extravasation (upper panel; white arrowheads) and Ter119 positive red blood cell extravasation (lower panel). Scale bar, 100 μm. **(H-l)** High-resolution confocal images of tip cell filopodia at the sprouting front (stained by IB4, marked by red dots) and respective quantification. Scale bar, 50 μm. (J) Confocal images EC proliferation (EdU^+^/ERG1^+^) at the angiogenic front. Scale bar, 50 μm. Error bars show s.d. of the mean. Statistical comparison by unpaired t-test with Welch’s correction. Number of analyzed animals (n) is indicated, ns = not significant, *p<0.05, **p<0.01, ***p<0.001.

PCR analysis of whole retinal lysates further confirmed the presence of a 380bp long *Srf*-exon 1 PCR product (*Srf-lx*) in *Srf^iMCKO^* mice, indicating successful recombination of the *Srf-flex1* allele. In contrast, in control mice, only a 1.34 kbp long PCR product, corresponding to the non-recombined LoxP flanked gene sequence of the *Srf-flex1* allele was amplified **(Online Figure I, D).**

Additional gene expression analysis of retinal MC populations sorted via fluorescence-activated cell sorting (FACS), showed efficient downregulation (over 99 %) of *Srf* expression **(Online Figure I, E)** in *Srf^iMCKO^* mice. Taken together, our conditional gene deletion approach allowed for successful and reliable deletion of *Srf* in MCs.

### SRF function in PCs is crucial for normal development of the retinal vasculature

In order to investigate the effects of mural *Srf* deletion on retinal angiogenesis, we analyzed *Srf^iMCKO^* and control retinas at P6. At this stage, the retinal vasculature is still actively sprouting and capillaries at the sprouting front become invested by PCs. At the same time, the more proximal vascular plexus is remodeling into a hierarchical network with arteries, arterioles, venules and veins. In *Srf^iMCKO^* retinas, radial vessel outgrowth was delayed and the vascularized area and number of capillary branches were decreased (**Figure *1*, C-E).** Increased levels of extravasated erythrocytes also suggested reduced barrier properties of the vasculature in *Srf^iMCKO^* retinas (**Figure *1,* F-G).** We further observed a decrease in the number of tip-cell filopodia and a severe dilation of blood vessels at the sprouting front (**Figure *1,* H-I).** In line with this, 5-ethynyl-2’-deoxyuridine (EdU) labeling of proliferating cells in combination with the EC specific nuclear marker ERG1 revealed an increase of EC proliferation in *Srf^iMCKO^* retinas (

**Figure *1,* J-K; Online Figure I, H).** Besides the abovementioned effects on the angiogenic sprouting front, the loss of *Srf* in MCs also affected remodeling of the proximal vascular plexus. We observed crossing of arteries and veins, which represents a microvascular abnormality termed nicking and is also observed in the human retina ^24^ (Online Figure I, F-G). We also noted abundant deposits of the basement membrane protein collagen IV without attached endothelial cells, so called empty matrix sleeves **(Figure 1, F),** which indicate excessive vascular pruning ^25^. These observations argue for defective remodeling and instability of nascent vessels in the proximal vascular plexus (**Figure *1,* F-G).**

### SRF mediates PC migration downstream of PDGFB via cytoskeletal re-arrangements

Since the vascular defects that we observed shared similarities with mouse models of postnatal PC depletion ^7,8,26^, we addressed PC recruitment and coverage in *Srf^iMCKO^* retinas. Immunostainings using antibodies against the PC markers NG2, DES and PDGFRB ^2^ revealed a 50.6 % reduction of PC coverage at the angiogenic sprouting front **(Figure 2, A-B; Online Figure II, A-B).** Interestingly, vessels at the central plexus showed only a 25.5 % reduction of PC coverage, suggesting that PC recruitment and migration along newly formed blood vessels was especially affected.

**Figure 2.**
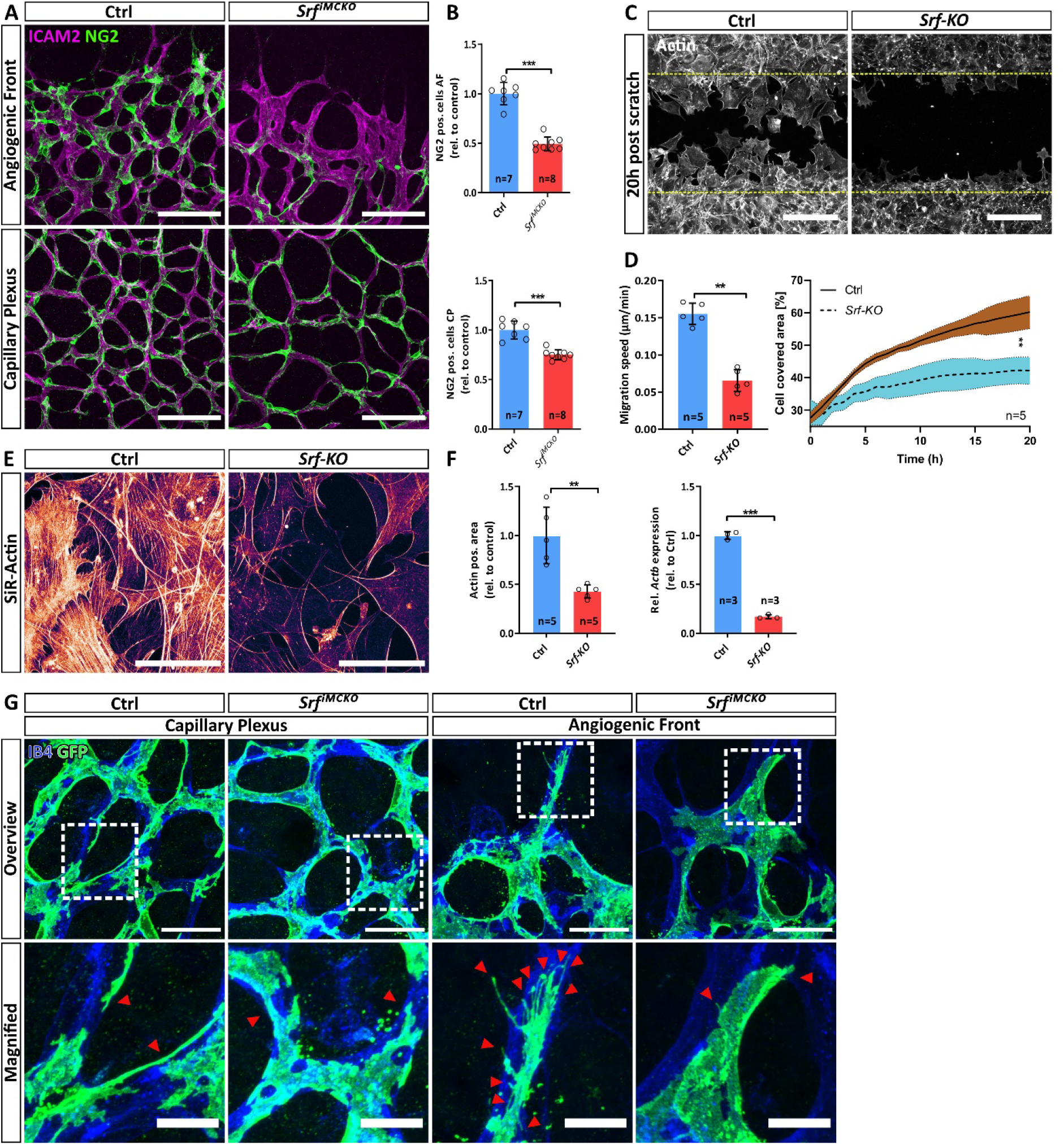
SRF controls cytoskeletal homeostasis and migration of pericytes (PCs). **(A)** Confocal image showing PC coverage (NG2,green) on vessels (ICAM2, magenta) at the angiogenic front (AF; upper panel) and the capillary plexus (CP; lower panel). Scale bar, 100 μm. **(B)** Respective quantification of PC coverage. **(C-D)** Confocal images of a 20h post scratch wound assay and respective quantifications using control and *Srf-KO* pBPCs, stained with the F-Actin staining dye SiR-Actin. Scale bar, 250 μm. **(E)** *Srf-KO* and control pBPC cultures stained with SiR-Actin. Note the reduction of F-actin in *Srf-KO* cells. Scale 100 μm. **(F)** Respective quantification of actin positive area and relative transcript levels of *Actb* determined by qPCR. (G) High resolution confocal images of PCs labeled by *mTmG* reporter expression (green) at the CP and the AF. ECs are stained with isolectinB4 (IB4). Red arrowheads point to tube like protrusions (in the capillary plexus) or filopodia (in the angiogenic front). Note the absence of filopodia as well as the shortened and flattened protrusions at *Srf^iMCKO^* PCs. Scale bars, 25 μm, 10 μm (magnified). Error bars show s.d. of the mean. Statistical comparison by Mann Whitney test. Number of independent animals/repetitions (n) is indicated. ns = not significant, *p<0.05, **p≤0.01, ***p≤0.001.

To specifically address if PC migration is compromised upon *Srf* deletion, we isolated primary PCs from the brain (pBPCs) of *Srf-flex1* and control mice and performed migration assays. In order to delete *Srf* in pBPC cultures, we exposed these cells to Tat-Cre, a membrane permeable version of the Cre recombinase. Subsequent qPCR analysis showed that this approach led to an over 99 % reduction of *Srf* mRNA and western blot analysis confirmed the loss of SRF protein **(Online Figure** II Error! Reference source not found., **C-D).** In scratch wound assays, *Srf* deleted pBPC cultures (hereafter referred to as *Srf-KO*) showed a significant reduction in collective cell migration, as well as in the speed of individual migrating cells **(Figure 2, C-D, Online Video** I). Likewise, we also observed reduced migration of *Srf-KO* cells in a trans-well assay **(Online Figure II, E-F).**

Since SRF is known to regulate cellular motility through transcriptional control of genes encoding regulators of actin dynamics in other cell types (reviewed in Olson & Nordheim, 2010) ^18^, we used the membrane permeable F-actin staining dye SiR-Actin ^27^, to visualize actin dynamics in living cells, and performed a series of time-laps experiments. These experiments revealed a substantial reduction of Factin in *Srf-KO* cells **(Figure 2, E-F).** In accordance, we also observed a significant reduction of beta actin gene (*Actb*) expression in *Srf-KO* cells **(Figure 2, F).** We further tested to which degree SRF mediated cytoskeletal re-arrangements are required downstream of PDGFB for PC migration. We therefore treated starved pBPCs with PDGFB and live imaged the changes in actin dynamics using SiR-Actin. Upon PDGFB stimulation, control pBPCs showed intensified actin dynamics and increased cell motility. In contrast, *Srf-KO* cells displayed almost no reaction to PDGFB stimulation **(Online Figure II, G; Online Video II).** Taken together, these results indicate that the observed migration defects of *Srf^iMCKO^* PCs are caused by the inability of the actin cytoskeleton to respond to the natural PDGFB gradient originating from endothelial tip cells.

In order to investigate PC morphology *in vivo,* we crossed the *Rosa26^mTmG^* reporter into the *Srf^iMCKO^* background and subsequently analyzed labeled PCs in *Pdgfrb-CreER^T2^::Roso26^mTmG^::Srf-flex1/Srf-flex1* (*Srf^iMCKO^*) and *Pdgfrb-CreER^T2^::Roso26^mTmG^::Srf-flex1/wt* littermate control retinas. In the absence of tamoxifen, the *Rosa26^mTmG^* reporter ubiquitously expresses tdTomato. However, upon tamoxifen induction, Cre mediated recombination leads to expression of membrane tagged eGFP which reliably outlines cell morphology **(Figure 2, G).** We found that control PCs at the capillary plexus attached tightly to the endothelium and extended numerous thin protrusions that connected PCs with each other. In contrast, *Srf^iMCKO^* PCs displayed an overall less ramified morphology and only formed short and stubby protrusions. At the angiogenic front, morphologic changes were even more pronounced. We noticed that control PCs often extended filopodia, which were oriented towards the angiogenic sprouting front, suggesting that PCs might use filopodia, similarly to ECs, for migration and to sense the PDGFB gradient **(Figure 2, G).** In contrast, SRF-deficient PCs had entirely lost the ability to form filopodia, appeared partially detached from ECs, and had adopted an abnormal cell morphology **(Figure 2, G).** The inability of *Srf^iMCKO^* PCs to form filopodia is in line with the actin remodeling defects that we observed in our *in vitro* experiments and consistent with SRF function in ECs where it also regulates filopodia formation ^28,29^. Taken together, our results suggest a crucial role for SRF in PC migration via regulation of actin dynamics.

### PDGFB signaling activates SRF via MRTF cofactors

The migration defects of SRF depleted PCs implied a direct role for SRF downstream of PDGFB signaling in PC migration. Most SRF-mediated motility responses have been reported to be regulated via RhoGTPase signaling. RhoGTPase activity stimulate F-actin polymerization, which depletes the cellular G-actin pool. Cytosolic G-actin can bind to myocardin-related transcription factors A and B (MRTF-A and MRTF-B) and thereby inhibiting nuclear translocation of MRTFs. Increased F-actin polymerization diminishes cytosolic G-actin levels, thereby enabling nuclear translocation of MRTF-A and MRTF-B and subsequent activation of SRF-directed transcription ^17^. To test if PDGFB also leads to SRF activation via MRTF cofactors, we took advantage of a 3T3 cell line expressing GFP-tagged MRTFA-protein ^30^. In order to observe potential MRTFA-GFP nuclear translocation upon PDGFB stimulation, we starved these cells overnight to cause MRTFA-GFP to predominantly localize in the cytoplasm **(Figure 3, A).** We subsequently imaged the starved cells using time-laps-microscopy and, after 15 minutes, stimulated with PDGFB **(Online Video III).** Strikingly, PDGFB stimulation led to a 3.5-fold increase in MRTFA-GFP nuclear translocation already within 5 min (T=20 minutes, **Figure 3, A-B)** and remained at high levels for another 5 min (T=25 minutes), before gradually shifting back to the cytoplasm. 35 minutes after stimulation (T=50 minutes, **Figure 3, A-B),** the nuclear MRTFA-GFP signal was reduced to 1.5-fold compared to pre-stimulation conditions and remained at this level for the rest of the experiment.

**Figure 3.**
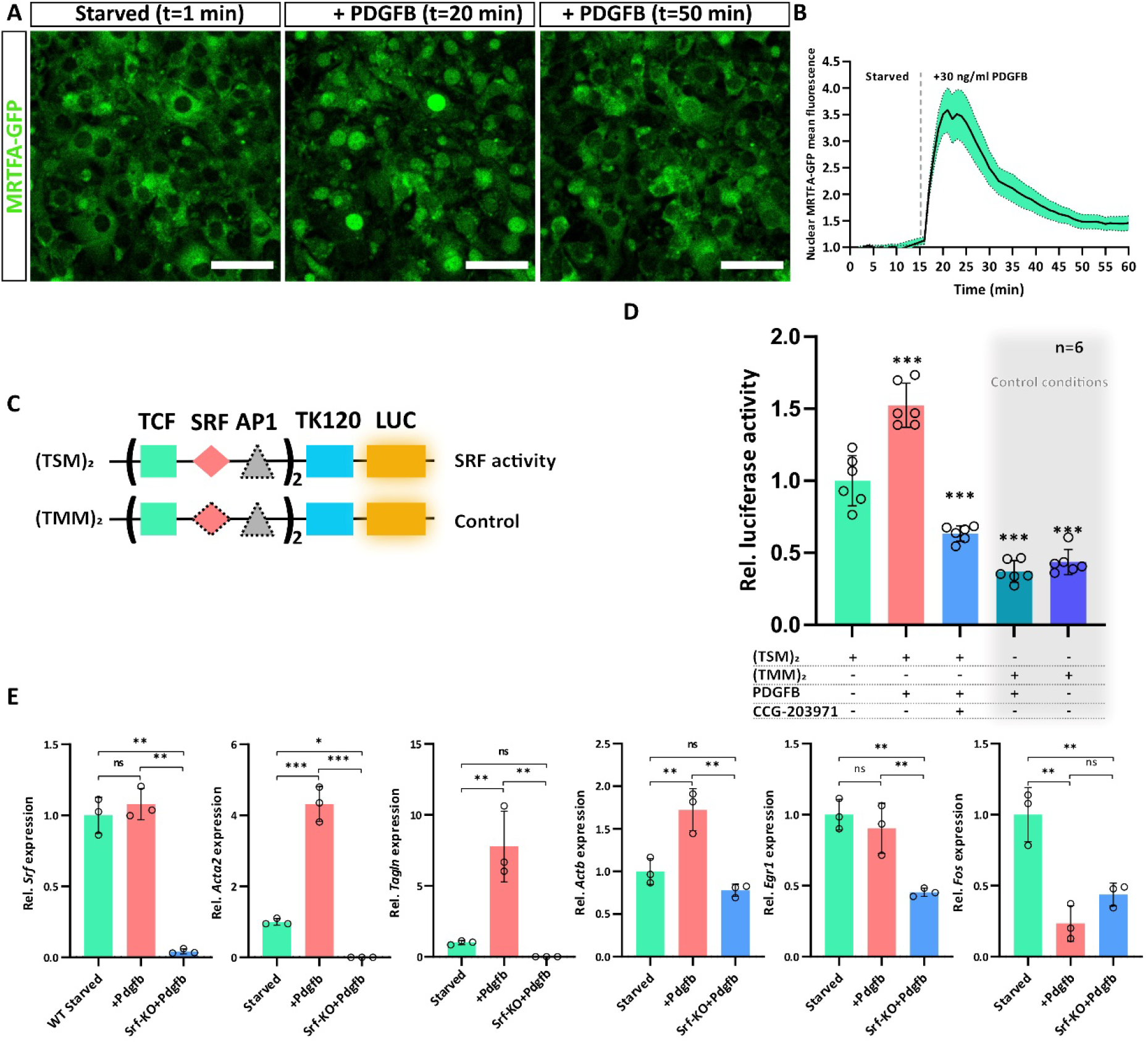
PDGFB signals towards SRF via MRTF cofactors. **(A)** Confocal live imaging of 3T3 cells stably transfected with MRTFA-GFP (green). The cells are shown before and after PDGFB stimulation. Time points are indicated. Scale bar, 50 μm. **(B)** Quantification of nuclear mean intensity of MRTFA-GFP signal over the time course of the experiment. In total, 29 cells were quantified. The experiment was carried out three times (n=3). Standard error of the mean (s.e.m.) is indicated in green. (C) Vector constructs used for luciferase assay, containing TCF, SRF and AP1 binding sites. Thymidine kinase minimal promoter (TK120) drives basal luciferase (LUC) expression. Symbols with dashed borders indicate mutated (unfunctional) binding sites. **(D)** Luciferase assay to verify SRF-driven luciferase activity upon PDGFB stimulation. Activity was modulated using vector constructs (TSM)_2_ and (TMM)_2_ as well as by using the MRTF inhibitor CCG-203971. (E) Gene expression analysis using qPCR of MRTF-SRF target genes (*Acta2, Tagln* and *Actb*) as well as TCF-SRF target genes (*Egr1* and *c-Fos*) in starved and PDGFB stimulated *Srf*-wild type (green and red bars) and *Srf-KO* (blue bars) pBPCs. Error bars indicate s.d. of the mean. Statistical comparison by one-way ANOVA (Tukey’s multiple comparison test). Number of independent repetitions is indicated. ns = not significant, *p<0.05, **p<0.01, ***p<0.001.

To investigate if PDGFB-induced nuclear MRTF translocation results in activation of SRF target genes, we performed luciferase-based reporter assays with promoter sequences containing functional (TSM)_2_ or mutated (TMM)_2_ SRF binding sites **(Figure 3, C)^28^.** PDGFB stimulation of 3T3 cells transiently transfected with the (TSM)_2_ reporter significantly increased basal luciferase activity **(Figure 3, D).** Addition of the MRTF inhibitor CCG-203971 abrogated the PDGFB-induced luciferase activation, demonstrating that PDGFB activates SRF mediated transcription primarily via MRTF co factors. The (TMM)_2_ construct served as negative control in those experiments, as SRF cannot bind to the mutated promoter sequence and thus is unable to activate the transcription of target genes. In accordance, PDGFB stimulation does not increase Luciferase activity in 3T3 cells transiently transfected with the (TMM)_2_ construct. To this end, qPCR analysis of PDGFB stimulated pBPCs further indicated a strong induction of the MRTF-SRF target genes *Actb, Acta2* and *Tagln* **(Figure 3, E).** Interestingly, we did not observe the induction of immediate early gene response genes *Egr1* and *c-Fos,* which are dependent on TCF-mediated SRF activation. Instead, TCF-SRF target genes were evenly downregulated. Taken together, our results strongly suggest that PDGFB-dependent SRF activation and transcription of target genes is mediated via MRTF co-factors.

### SRF is a key determinant of pathologic PC activation

Recent work of Dubrac *et al.* (2018) has shown that, under certain pathologic conditions, PCs can acquire disease promoting properties ^31^. Using the oxygen induced retinopathy (OIR) mouse model, it was shown that excessive PDGFB-PDGFRB signaling leads to pathological activation of PCs, which promotes the formation of neovascular tufts (NVT). NVTs are clustered capillary loops, which show excessive EC proliferation and extravasation of RBCs. During OIR, PCs undergo a pathological switch accompanied by upregulation of SMGs, which is characterized by strong expression of αSMA. Since our results suggested that SRF regulates both PC migration and expression of αSMA, we hypothesized that SRF might be a driving force of pathological PC activation during ischemic conditions. To address this hypothesis, we performed OIR experiments and kept P7 *Srf-flex1::Pdgfrb-CreER^T2^* and control pups for 5 days under hyperoxic conditions, which led to vaso-obliteration in the primary vascular plexus **(Figure 4, A).** We then returned the mice to normal oxygen conditions (21 % O_2_), which provoked a strong hypoxic response and resulting pathological revascularization and NVT formation. During the revascularization phase, we applied tamoxifen to the pups (P12-P14) to induce *Srf^iMCKO^* and analyzed the impact on NVT development at P17.

**Figure 4.**
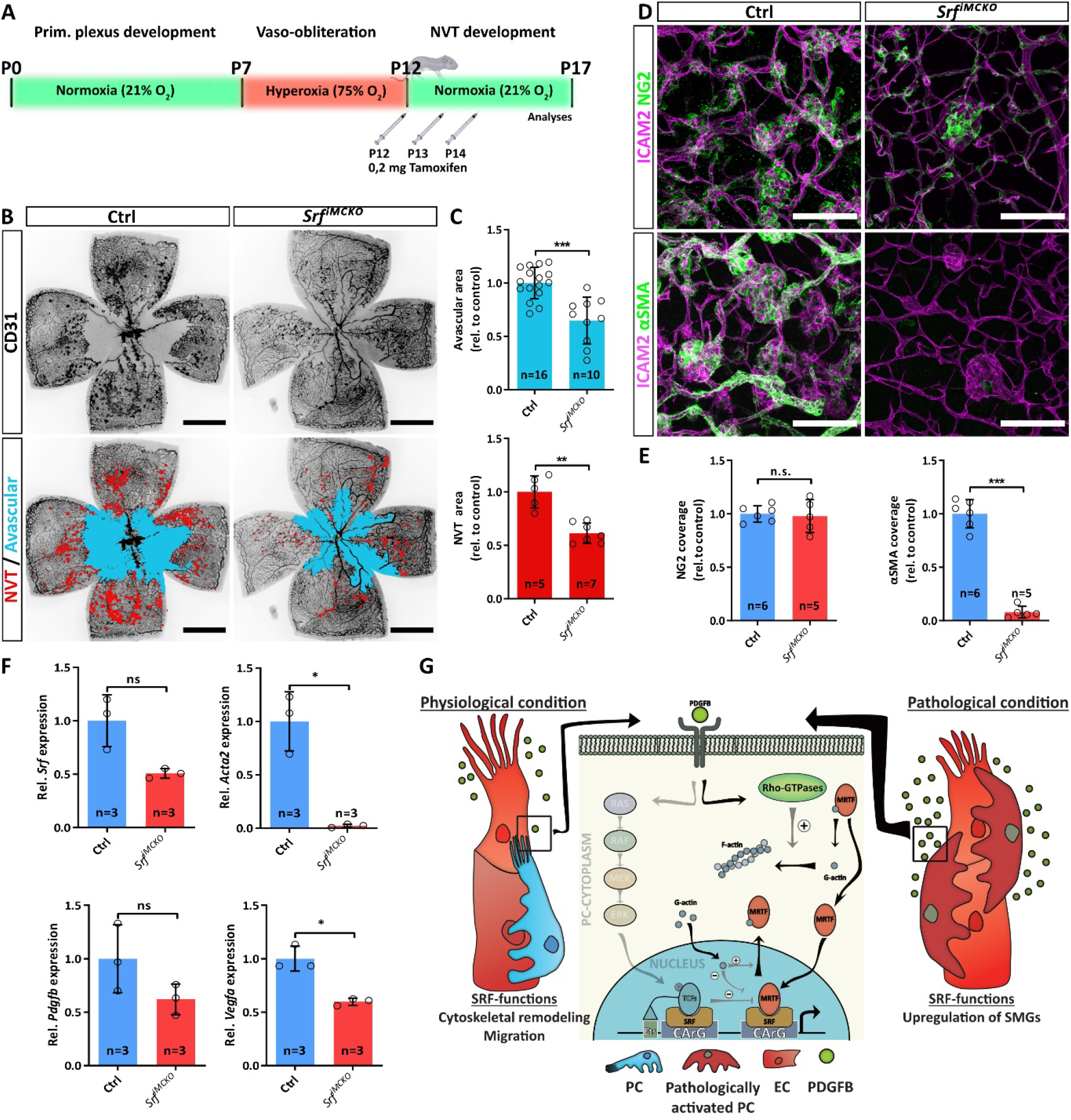
SRF is a key determinant of pathological pericyte (PC) activation. **(A)** Experimental outline of oxygen induced retinopathy (OIR) experiments (P indicates postnatal days). (B) Epifluorescence overview images of *Srf^iMCKO^* and control OIR retinas. The avascular (blue) and neovascular tuft (NVT) area (red) are highlighted in the lower panel. Note the reduction of avascular and NVT area in *Srf^iMCKO^* retinas. Scale bars, 1 mm. (C) Quantification of avascular (upper graph) and NVT area (lower graph). (D) High resolution confocal images of NVTs in control and *Srf^iMCKO^* OIR retinas stained by NG2 (green, first row), αSMA (green, second row) and the endothelial cell marker ICAM2 (magenta, first row). Scale bar, 100 μm. (E) Quantification of PC coverage by NG2 staining as well as pathologically activated PCs by αSMA staining. (F) Quantification of relative transcript levels of *Srf, Acta2, Pdgfb* and *Vegfa* determined by qPCR from whole OIR retina lysates. (G) Mechanistic model of SRF guided PC functions under physiological and pathological conditions. Error bars show s.d. of mean. Statistical analysis by unpaired t-test with Welch’s correction. Number of analyzed animals (n) is indicated. ns = not significant, *p<0.05, **p<0.01, ***p<0.001.

Stainings with the endothelial marker CD31 revealed a significant reduction of NVT development by 38.5 % (±9.5 %; p=0.0025; **Figure 4, B-C)** and exhibited an improved revascularization, evident as a reduction of the avascular area by 35.2 % (±21.2 %; p=0.0005) in *Srf^iMCKO^* OIR retinas. Co-stainings for NG2 and desmin confirmed the presence of PCs on NVTs both in control and *Srf^iMCKO^* OIR retinas **(Figure 4, D; Online Figure III, A-B).** While in control retinas pericytes on NVTs displayed the characteristic upregulation of αSMA **(Figure 4, D),** the αSMA staining in *Srf^iMCKO^* OIR retinas was reduced by 92% **(Figure 4, D-E).** We further confirmed by qPCR analysis that mRNA expression of *Acta2,* the gene encoding αSMA, was almost completely lost in *Srf^iMCKO^* OIR retinal lysates **(Figure 4, F),** which strongly suggests that pathologic activation of PCs was diminished. In addition, we observed that mRNA levels of *Vegfa,* the main angiogenic driver, were also significantly reduced **(Figure 4, F).** This, in turn, could explain the reduction in NVT formation and decrease in avascular area.

Taken together, our results show that SRF is a necessary player in the pathogenic activation of PCs during OIR and that αSMA upregulation, a common finding in several vascular conditions, relies on the activation of this transcription factor. The relevance of SRF for the phenotypic switch of PCs during OIR makes it a potential target to prevent pathologic activation of PCs although its role during physiologic angiogenesis **(Figure 4, G)** should not be overlooked.

### SRF deletion in MCs triggers the formation of arteriovenous malformations

Since in *Srf^iMCKO^* mice, *Srf* is not only deleted in PCs but also in vSMCs, we wanted to study its requirement for vSMC development and function in further detail. In order to visualize vSMCs during early retinal development, we performed immunohistochemical staining for αSMA and NG2 at P6. In control retinas, αSMA strongly highlighted part of the contractile machinery located in the cytoplasm of vSMCs, whereas membrane-bound NG2 staining outlined the cell shape. Remarkably, in vSMC of *Srf^iMCKO^* retinas, the αSMA signal was lost **(Online Figure IV, A).** qPCR analysis of whole retina lysates from *Srf-KO* pBPCs confirmed a dramatic drop of *Acta2* gene expression **(Online Figure IV, B).** Despite the complete absence of the αSMA signal, NG2 staining still marked the vSMC population on arteries **(Online Figure IV, A-B),** indicating that vSMCs were present but had lost *Acta2* expression. An in-depth analysis of NG2-positive cell coverage on arteries indicated only a slight reduction of vSMCs in *Srf^iMCKO^* retinas **(Online Figure IV, B).** These results strongly argued that SRF is strictly required for αSMA expression in arterial vSMCs.

To investigate the impact of *Srf* deletion during vascular remodeling, we analyzed retinas at P12. At this stage, blood vessels sprout perpendicular form the primary retinal plexus in order to form the deep plexus^32^. The tissue undergoes extensive remodeling and both arteries and veins progressively mature. vSMCs become increasingly important as the blood pressure in arteries increases, and consequently contractile functions are crucial to regulate the vascular tone. At this stage, control retinas displayed a stereotypical vascular pattern, with hierarchical organized blood vessels and a clear arterial-venous zonation, implicating that blood flow is channeled from arteries into arterioles, capillaries, venules and subsequently into veins. In contrast, we observed severe patterning defects in *Srf^iMCKO^* retinas ( **Figure 5, A),** in which both arteries and veins were significantly dilated ( **Figure 5, B-C).** Moreover, arteries often formed direct connections to the veins, forming arteriovenous (AV)-shunts (**Figure 5, A).** The venous portion of the AV-shunt was often severely dilated and misshaped. This phenotype was highly penetrant, since we observed at least one AV-shunt in every analyzed P12 retina. In addition, we noticed that the deep vascular plexus was considerably under-developed, as illustrated by a reduced vascular area and a decreased number of vessel branch points (**Figure 5, D-E).** Remarkably, deep plexus capillaries were completely absent directly underneath the AV-shunts (**Figure 5, A).**

**Figure 5.**
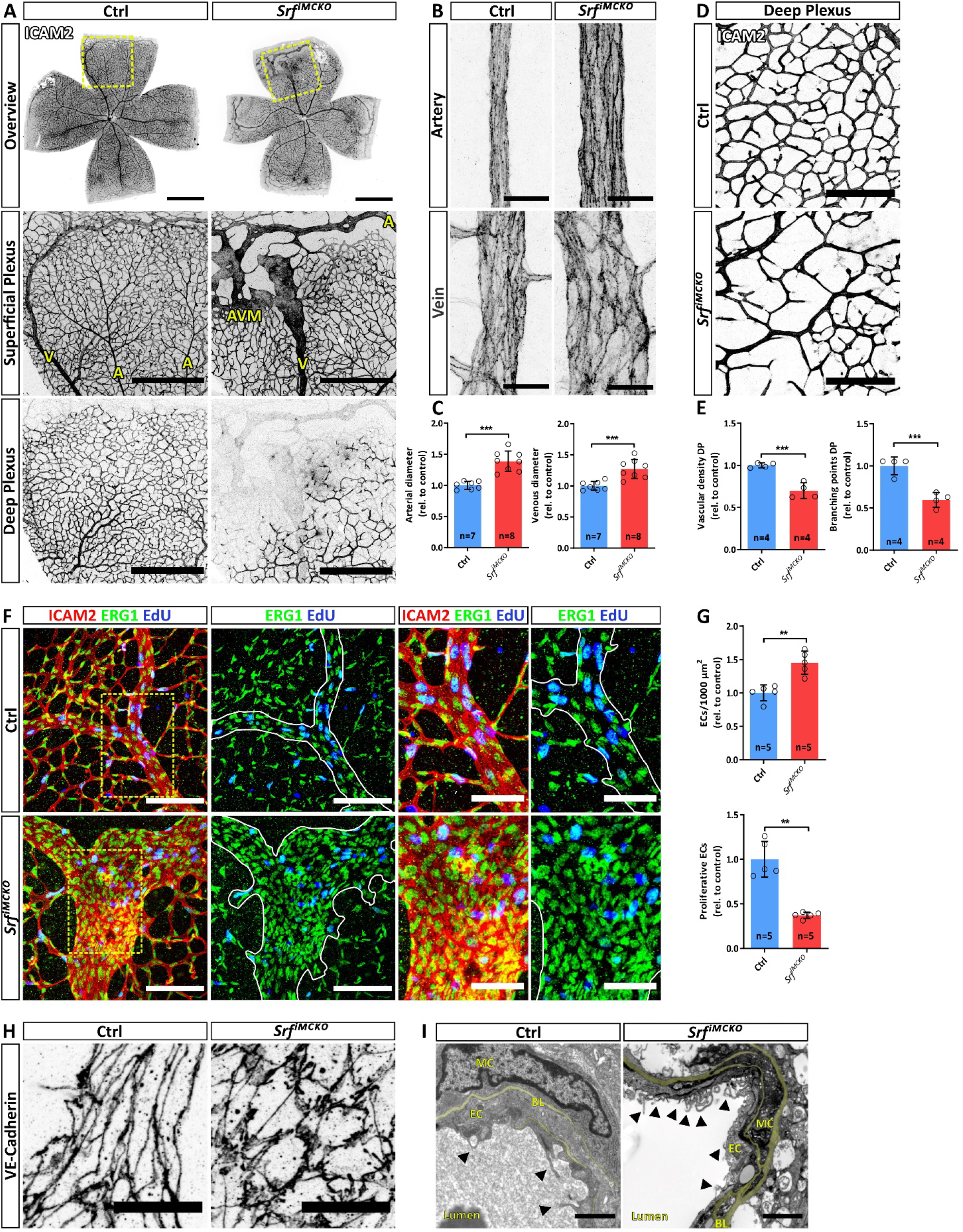
SRF-deficient vascular smooth muscle cells (vSMCs) trigger the formation of arteriovenous (AV)-shunts. **(A)** Epifluorescence overview (upper row) and confocal images (lower panels) of P12 control and *Srf^iMCKO^* retinal whole mounts, stained using the endothelial marker ICAM2. The yellow dashed squares indicate magnified regions of superficial and deep plexus (middle and lower columns). Arteries (A), veins (V) and the arteriovenous malformation (AVM) are annotated in yellow letters. Scale bar: 1 mm (upper row), 500 μm (middle and lower row) (B) Confocal, images of representative arteries and veins indicating the dilation of both vessel types in *Srf^iMCKO^* retinas. **(C)** Quantification of artery and vein diameters. **(D)** Representative confocal image of the deep plexus. (E) Quantification of the vascular density. Scale bar, 150 μm. (F) AV-shunts and control veins, showing proliferating cells (EdU, blue), blood vessels (ICAM2, red) and endothelial nuclei (ERG1, green). White lines define the border of malformations or respective control positions, and outline the vessel shape (second and forth column). Images in the third and fourth columns show magnification of the yellow dashed squares of the panels in the first column. Scale bar, 100 μm (left) and 50 μm (right). (G) Quantification of the EC density (ERG1+ counts/ICAM2+ area) and EC proliferation (ERG1^+^ counts/ERG1^+^+EdU^+^ counts). **(H)** High resolution confocal image of the junctional EC-specific protein VE-Cadherin, visualizing the shape of endothelial cells. Scale bar, 20 μm. (I) Electron microscopy images of control veins and malformed *Srf^iMCKO^* veins, visualizing ultrastructural changes. Black arrowheads pointing to EC membrane invaginations. Note also that the basal lamina (BL, pseudocolored in yellow) is thickened in *Srf^iMCKO^.* Scale bar, 1 μm. Error bars indicate s.d. of the mean. Statistical comparison by unpaired t-test with Welch’s correction. Number of analyzed animals (n) is indicated. ns = not significant, *p≤0.05, **p≤0.01, ***p≤0.001

The AV-shunts in *Srf^iMCKO^* retinas morphologically resembled vascular malformations characteristic of mouse models of hereditary hemorrhagic telangiectasia (HHT), a disease caused by mutations in the TGFß pathway^33^. In HHT mouse models, it has been reported that ECs that contribute to shunt-formation proliferate at higher rates than neighboring ECs ^34,35^. To clarify if a similar mechanism could explain AV-shunt formation in *Srf^iMCKO^* retinas, we performed *in vivo* proliferation assay using 5-ethynyl-2’-deoxyuridine EdU in combination with the nuclear EC marker ERG1. Whereas ERG1 staining revealed that EC density was locally increased in the malformed areas, EC proliferation (ERG1^+^ + EdU^+^ counts/ERG1^+^ counts) was markedly reduced in *Srf^iMCKO^* retinas (**Figure 5, F-G).** Immunostainings with the junctional marker VE-cadherin further demonstrated that the shape of EC in the malformed regions was severely affected. While ECs on veins in control retinas were elongated and showed a spindle-like morphology with straight adherence junctions, ECs at AV-shunts had a more round morphology, were less elongated with irregular adherence junctions that appeared to be partially overlapping and had a zick-zack morphology (**Figure 5, H).** Furthermore, transmission electron microscopy (TEM) of AV-shunt regions revealed enlarged basallaminae and intraluminal membrane invaginations originating from ECs (**Figure 5, I**). Interestingly, intraluminal membrane invaginations have also been described in *Pdgfb* mutant mice, in which PC recruitment is defective^36^.

### SRF is critical for the expression of contractile genes in vSMCs

While, overall, we only observed a marginal difference in MC coverage between control and *Srf^iMCKO^* retinas **(Figure 6, A; Online Figure V, A-D),** we noticed, in AV-shunt areas, that the venous shunt-portions showed an increased vSMC coverage, while arterial shunt-portions were often deprived of vSMCs **(Figure 6, A; Online Figure V, C-D).** A 3D segmentation analyses of AV-shunt regions showed striking morphological alterations of vSMCs **(Figure 6, B)** and high-resolution imaging further revealed changes in the intracellular organization of the cytoskeletal protein desmin **(Online Figure V, E).** Taken together with the complete absence of αSMA **(Figure 6, C-D)** these results suggested that the contractile ability of vSMCs is compromised in *Srf^iMCKO^* retinas. To further investigate a potential reduction of vSMC contractility, we determined the expression levels of genes, encoding typical contractile proteins in SMCs ^29, 30^ and found that *Acta2, Mhy11* and *Tagln* were strongly downregulated in whole retinal lysates of *Srf^iMCKO^* mice **(Online Figure VI, A).** We further used pBPCs cultures to study the expression of contractile genes. In control pBPCs cultures, the addition of serum leads to a strongly induction of SMGs, such as *Acta2, Mhy11* and *Tagln,* whereas in *Srf-KO* cells the expression of those genes was almost completely lost in any tested condition **(Online Figure VI, B).** Altogether, these results show that SRF is indispensable for the expression of genes responsible for the contractile abilities of vSMCs.

**Figure 6.**
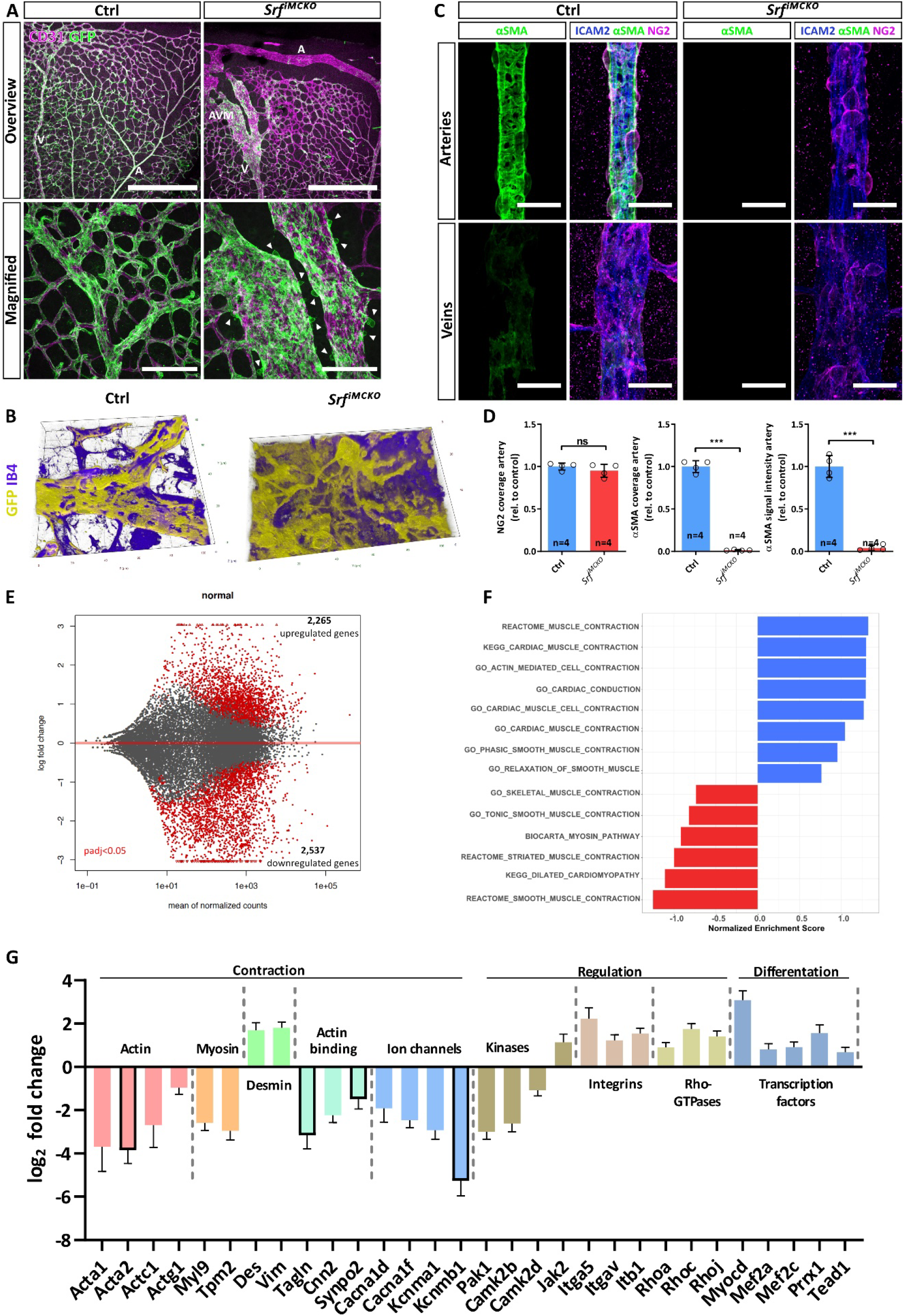
Dysregulated expression of contractile genes in *SRF^iMCKO^* mural cells (MCs). **(A)** Confocal images of retinal vasculatures genetically labeled with the *Rosa26^mTmG^* reporter for MCs (GFP, green) and co-stained for CD31 (ECs, blue). Arteries (A), veins (V) and arterio-venous malformation (AVM) are highlighted with white letters. Note MC free area of arteries at the periphery in the *Srf^iMCKO^* retina. White arrowheads point at MCs, which round up and appeare to detach from the endothelium in *Srf^iMCKO^* animals. Scale bars, 500 μm (top panel) 100 μm (lower panel). (B) 3D rendering of confocal images showing morphological abnormalities of MCs in AV-shunts of *Srf^iMCKO^* retinas and corresponding venous areas in control retinas. **(C)** Confocal images of arteries and veins of *Srf^iMCKO^* and respective control retinas stained for αSMA (green), ICAM2 (blue) and NG2 (magenta). Note the loss of αSMA signal in arterial and venous MCs of *Srf^iMCKO^* retinas despite the presence of NG2. Scale bar, 25 μm. (D) Quantification of aSMC coverage determined by NG2-/ICAM2 area, αSMA-/ICAM2 area and αSMA signal intensity on arteries. (E) Vulcano plot displaying RNA-Seq detected genes according to the degree of dysregulation (log2 fold change). Red dots indicate significantly (p<0.05) dysregulated genes. (F) Gene set enrichment analysis (GSEA) of RNA-Seq dataset of *Srf^iMCKO^* and control MCs. Positive normalized enrichment scores (NES) indicate pathways containing downregulated genes whereas negative NES indicate pathways containing genes which are upregulated. Representative summary of tested contraction-related gene sets within the GO, Reactome and Biocarta databases. **(G)** Gene expression log2 fold changes of selected genes identified by the RNA-Seq methodology and subsequent GSEA. Listed genes are essential contributors to smooth muscle cell (SMC) contraction and were functionally grouped (headings). Framed bars indicate frequently used SMC markers. Error bars indicate the standard error of the mean (s.e.m.). All presented genes were significantly dysregulated (p<0.05). Statistical comparison by unpaired t-test with Welch’s correction. Number of analyzed animals (n) is indicated, ns = not significant, *p≤0.05, **p≤0.01, ***p≤0.001

In order to characterize the transcriptional changes that result from SRF deletion in MC in further detail, we utilized an RNA sequencing (RNA-Seq) approach. We induced control and *Srf^iMCKO^* mice with tamoxifen from P1 to P3 and FAC-sorted PDGFRB^+^ MCs from retinas at P12. We subsequently isolated RNA from those cells and sequenced in triplicates. Expression analysis confirmed a high enrichment of mural specific genes such as *Pdgfrb, Rgs5* and *Notch3,* whereas genes typically expressed in other cell types such as ECs (*Pecam1, Cdh5*) or astrocytes (*Gfap*) were underrepresented, which suggested that sufficiently pure MC fractions had been isolated and sequenced **(Online Figure VI, C).** Principal component analysis (PCA) of all sequenced datasets showed a sufficient reproducibility between samples within one group and a strong difference between the control and the *Srf^iMCKO^* group **(Online Figure VI,** D).

Differential gene expression analysis using a false discovery rate (FDR)-adjusted p-value < 0.05 and an absolute log_2_ fold change (fc) > 0.5 identified 2,265 upregulated and 2,537 downregulated genes **(Figure *6*, E).** Notably, we identified 517 differential expressed genes (DEGs; 350 up/167 down) which are under control of SRF ^37,38^. The expression of the majority of these DEGs (375) is controlled by the MRTF-SRF signaling axis **(Online Figure VI, E).** Further, gene set enrichment analysis (GSEA) using the Kyoto Encyclopedia of Genes and Genomes (KEGG) and the Gene Ontology (GO) databank resulted especially in the identification of pathways linked to contractile functions of myocytes and SMCs **(Figure 6, F).** Importantly, the GSEA identified numerous DEGs crucial for smooth muscle contraction **(Figure 6, G),** and the downregulation of Acta2 and *Tagln* in *Srf^iMCKO^* MCs could be confirmed. Strikingly, we also observed a strong downregulation of numerous structural elements of the contractile apparatus such as myosin light chain 9 (*Myl9*; log_2_fc: −2.59 ±0.36), tropomyosin beta chain (Tpm2; log_2_fc:-2.96 ±0.42) and several members of the actin family such as *Acta1* (log_2_fc: −3.68 ±1.14), *Actc1* (log_2_fc: −2.69 ±1.03) and *Actg1* (log_2_fc: −0.97 ±0.31). Of note, multiple ion channel proteins important for Ca^2+^ release into the cytoplasm, which triggers vSMC contraction, were markedly downregulated (*Cacna1d,* log_2_fc: −1.91 ±0.64; *Cacna1f,* log_2_fc: −2.46 ±0.34; *Kcnma1,* log_2_fc: −2.92 ±0.42 and *Kcnmb1,* log_2_fc: −5.27 ±0.7). Furthermore, we noticed a partial downregulation of kinases which are involved in the regulation of the vascular tone (Camk2b; log_2_fc: −2.61 ±0.39 and Camk2d 1.09 ±0.26 and Pak1; log_2_fc:-3.0 ±0.34). Conversely, several integrins, Rho-GTPases and transcriptional elements were found to be upregulated in *Srf^iMCKO^* MCs. Most notably the SRF-cofactor myocardin (*Myocd,* log_2_fc: 3.08 ±0.43), which binds to and activates SRF and thereby leads to induction of muscle-specific gene expression^39^. The upregulation of those genes might reflect a compensatory mechanism aimed to limit the consequences that the loss of SRF caused for mural cell function.

### Loss of vascular tone in the absence of mural SRF

Our transcriptomic analyses of MCs, isolated from P12 *Srf^iMCKO^* and control retinas revealed a conspicuous mis-regulation of genes involved in the regulation of the vascular tone. In order to explore the biological significance of these results, we investigated the retinal vascular morphology of control and *Srf^iMCKO^* mice at 3, 4 and 8 weeks of age *in vivo* via scanning laser ophthalmoscopy (SLO) and optical coherence tomography (OCT). SLO confirmed a substantial enlargement of arterial and venous vessels in all examined *Srf^iMCKO^* retinas. Moreover we were able to classify the individual vascular alterations into three different degrees of severity, ranging from mild and intermediate to severe **(Online Figure VII, A-B).**

Strikingly, there were also major abnormalities in the dynamic movement of vessel walls associated with blood pulsations in *Srf^iMCKO^* retinas. Whereas the vessel wall of arteries in control retinas showed relatively scant movements, we observed exceptionally strong pulsating motion in the arterial wall of *Srf^iMCKO^* animals **(Figure 7, A-C; Online Video IV).** This finding is in strong support of a hemodynamically relevant loss of the vascular tone. Moreover, in our longitudinal study, we further observed that arterial and venous vessels gradually enlarged during the course of the experiment. This effect was quantified by repetitive size assessment of identical vessels at 3, 4 and 8 weeks, respectively **(Figure 7, D).** In the most severe cases, we observed that venous vessels ruptured **(Figure 7, D),** which further supports the hypothesis that vSMCs had lost their ability to modulate blood pressure.

**Figure 7.**
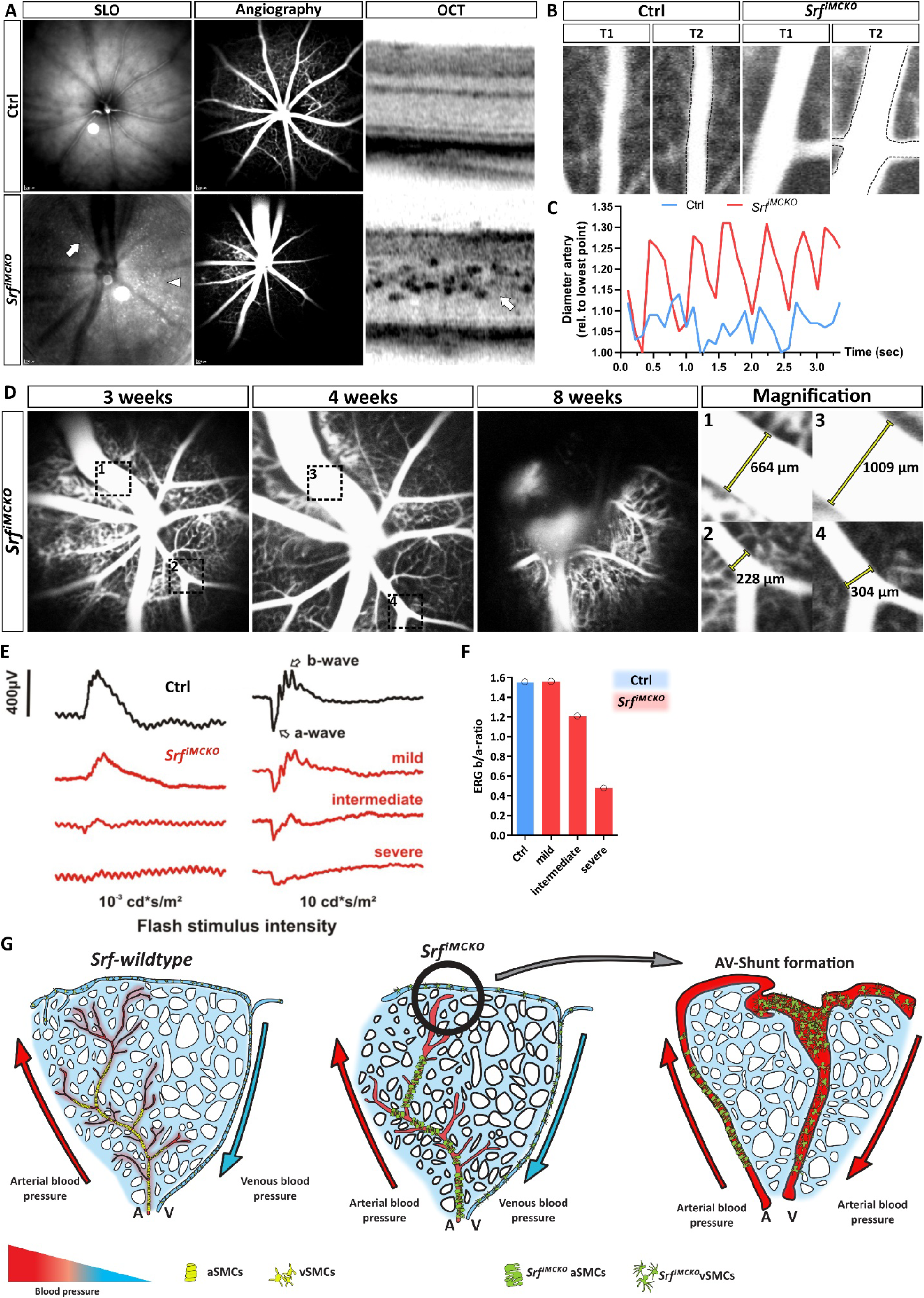
*Srf^iMCKO^* vascular smooth muscle cells (vSMCs) lose their ability to maintain the vascular tone. **(A)** Scanning laser ophthalmoscopy (SLO) and optical coherence tomography (OCT) live *in vivo* imaging of 4 weeks old control and *Srf^iMCKO^* animal. Native SLO and Indocyanine green (ICG) perfused angiography highlighting vessel structure and perfusion whereas OCT imaging highlighting optical section of the retina. (B) Representative control and *Srf^iMCKO^* artery at two timepoints (T1 and T2) indicating vessel movements. The dashed black line indicates Tl. **(C)** Respective measurement of arterial vessel diameter shown in (B). **(D)** Serial angiography imaging of the same *Srf^iMCKO^* eye at 3, 4 and 8 weeks. Note the rupture of the venous vessel at 8 weeks. The right panel shows magnification of the numbered and dashed black boxes shown in the first two panels. Venous (1 and 3) and arterial (2 and 4) diameters are annotated. **(E)** Examples of *in vivo* electroretinography (ERG) data from the scotopic single flash intensity series, obtained from a control and 3 differently affected mutant animals at the age of 4 weeks. Note the reduced overall size of responses in the mutants, together with a relatively strong reduction of the b-wave component in comparison to the a-wave component. (F) Quantification of the ERG b/a-ratio for the results shown in (E) indicating increasing retinal hypoxia and a growing functional deficit of vision (G) Model depicting how the loss of vascular tone could trigger the development of arteriovenous (AV)-shunt formation in *Srf^iMCKO^* mutant mice.

To investigate potential perfusion problems that could result from a loss of vascular tone, we analyzed *Srf^iMCKO^* and control retinas with SLO angiography, for which we used indocyanine green (ICG) as a contrast agent. These results revealed profound alterations of the capillaries in *Srf^iMCKO^* retinas, which are suggestive of associated perfusion defects **(Figure 7, A; Online Figure VII, A).** The seemingly reduced capillary perfusion likely has detrimental effects on the retinal tissue, as oxygenation and nutrient supply are expected to be severely decreased. Consequently, we performed an assessment of retinal function via electroretinography (ERG). The electroretinogram is a measure of retinal sum potentials triggered by a brief light stimulus and mainly comprises the transient activity of photoreceptors and bipolar cells. A series of light flashes of increasing intensity is commonly employed to investigate the integrity of the visual system at the level of mainly the outer retina ^40^. A typical finding in the dark adapted (scotopic) ERG in case of retinal hypoxia is a discrepancy between the initial negative wave (the a-wave) and the subsequent positive wave (the b-wave), leading to a waveform shape called ‘negative ERG’.

Indeed we clearly observed ‘negative’ scotopic ERG in eyes of *Srf^iMCKO^* animals as well as reduced b/a-wave ratios **(Figure 7, E-F),** indicating reduced retinal oxygenation, most likely as a result of defective capillary perfusion. The grade of severity of vascular alterations was well correlated with the scotopic ERG measurements **(Figure 7, E-F; Online Figure VII, A).**

To analyze if SRF is also needed for mural cells function in the mature vasculature, we induced its deletion in adult mice. We injected 8-week old *Srf-flex1::Pdgfrb-CreER^T2^* and littermate control mice with tamoxifen on 5 consecutive days **(Online Figure VIII, A)** and analyzed retinas after 2 and 12 months respectively. After 2 months, the retinas showed no obvious vascular phenotype (data not shown). However, after 1 year, we observed a significant dilation of arteries and veins **(Online Figure VIII, B-C).** Co-immunostainings for desmin and αSMA did not indicate a substantial loss of vSMC coverage on arteries or veins, but revealed a marked reduction of αSMA expression, especially on veins **(Online Figure VIII, D-E).** This suggests that SRF deletion in adult mice leads to a reduction in contractile protein expression rather than a loss of vSMCs. In support of this hypothesis, we also observed a reduced expression of *Acta2* and *Tagln* in whole brain lysates of in *Srf^iMCKO^* mice **(Online Figure VIII, F).** In contrast, we did not observe changes in the capillary network of those mice and conclude that SRF is dispensable in adult retinal PCs.

## 4. Discussion

Our understanding of MC function has increased considerably in the past decade. It is now well established that PCs play a key role in maintaining the integrity of the BBB and that their dysfunction contributes to the progression of numerous diseases (reviewed in ^41–43^). Yet, despite recent advances, many aspects of PC biology still remain poorly understood.

During angiogenesis, PCs are recruited to sprouting blood vessels via PDGFB/PDGFRB signaling ^11,44^. PDGFB, which is produced and secreted by tip cells is retained in the extracellular matrix (ECM) of new vessel sprouts via its retention motive ^45^. PCs, which in turn express PDGFRB, sense the PDGFB tissue gradient and co-migrate along the nascent vessels sprouts ^46^. NCK1 and NCK2 adaptor proteins have been proposed to mediate PDGFB dependent PDGFRB phosphorylation ^31^. However, which signaling molecules are activated downstream of PDGFRB during PC recruitment and how those molecules regulate the cytoskeleton in order to mediate cell motility has not been well characterized. Here, we demonstrate that PDGFB/PDGFRB signaling triggers translocation of MRTF co-activators to the nucleus, where they associate with the SRF transcription factor and activate expression of a specific gene set that subsequently regulate PC migration. Interestingly, MRTF-driven activation of SRF has previously been reported in response to PDGFRA signaling in mesenchymal cells during craniofacial development ^47^, suggesting that MRTF/SRF activation might be a conserved feature downstream of PDGFRs. Our data suggests that SRF is a key regulator of cytoskeletal functions in PCs, as its deletion (*Srf^iMCKO^*) led to severe cytoskeletal and migratory defects *in vivo.* As a consequence, SRF-depleted PCs were unable to fully populate the retinal vasculature, which resulted in a reduced PC coverage, especially at the sprouting front, and caused vessel dilation as well as reduced barrier properties.

Recent studies have highlighted that PCs are not only crucial for normal vascular development and for maintaining BBB barrier properties in the adult vasculature but can also acquire disease promoting properties. Examples are the formation of vascular malformations as a consequence of RBP-J deletion in PCs and their role as promoters of NVT formation in ischemic retinopathy^31,48^.

Here we demonstrate that the pathological features of PC activation in ischemic retinopathy are mediated by SRF, which regulates PC migration downstream of PDGFRB signaling and activates the expression of SMGs. Accordingly, OIR experiments in *Srf^iMCKO^* mice showed reduced NVT formation, and improved revascularization. Remarkably, the pathological αSMA expression in PCs was completely prevented.

In this regard, it is interesting to note that pathological activation of PCs shares certain similarities with fibrotic reactions in which SMGs are expressed at high levels and excessive amounts of ECM proteins are deposited ^49^. The fibrotic reaction is regulated, at least in part by the actin-MRTF-SRF axis^50^, and recently developed small molecule inhibitors that target MRTF function are promising candidates for the treatment of fibrosis ^51^.

Besides its crucial role in PCs, SRF also plays essential roles in vSMCs, where it regulates the expression of SMGs. These genes typically contain a CArG (-cis) element that serves as a SRF binding side ^37,52^ and, in part, encoded proteins that enable vSMCs to constrict and thereby to increase vascular resistance ^53,54^. Through the modulation of vascular resistance, vSMCs are able to regulate blood flow in order to satisfy the local demands for oxygen and nutrients ^55^. This implies, that a vessel branch that experiences a stochastic increase in flow compared to its neighboring branch must be able to counterbalance that increased flow rate to ensure an equal blood distribution. This is attained by an increase in resistance in the affected branch due to vasoconstriction, which naturally leads to an increased flow in the neighboring branches, where resistance is lower ^56,57^. This property of vSMCs has been termed the myogenic response^58^. In *Srf^iMCKO^* mice, vSMCs fail to express typical SMGs and are no longer able to mediate the myogenic response. Consequently, flow redistribution cannot be achieved and initial stochastic changes in local blood flow cannot be adequately redirected. We propose, that, as a consequence, some branches develop into AV-shunts that funnel a proportion of the blood directly to the venous circulation **(Figure 7, E).** This relives the pressure from surrounding vessels and ensures a certain functionality of the retinal vasculature. These shunts form, where one would expect, in the retinal periphery, where the distance between arteries and veins is the shortest. Nevertheless, electrophysiological measurements revealed that retinal function and thus, vision, is severely impaired by the reduced perfusion.

In this context it is interesting to note that similar roles of SRF have been reported in visceral SMCs, where *Srf* deletion led to impaired contraction and thus to severe dilation of the intestinal tract^19,22^. The fully developed retinal vasculature seems to be more robust to changes. Mural *Srf* deletion in adult mice did not lead to AV-shunt formation which is likely attributed to the low plasticity of fully matured blood vessels. In this context, it is worth noting that adult deletion of SRF did not result in a complete loss of αSMA in arterial SMCs. It is thus possible, that the remaining αSMA protein levels maintain a sufficient degree of contractile function and that, because of this, AV-shunts did not form. However, the diameter of arteries and veins significantly increased in aged *Srf^iMCKO^* mice, and we observe a significant reduction of contractile proteins. In contrast, PCs at the capillaries seemed unaffected, suggesting that SRF is dispensable for PC homeostasis.

The finding that defective MC function can trigger the formation on AVMs, might be of broader medical relevance. AVMs are hallmarks of HHT, a human disease caused by autosomal dominant mutations in genes of the TGF-β signaling pathway, in particular ENG or ACVRL1^33^. In HHT, AVMs commonly form in the nose, lungs, brain or the liver and affected individuals often suffer from nasal and gastrointestinal bleedings. Rarely occurring AVMs in the CNS can even be live threatening^59^. Thus far, MCs have not been directly implicated to trigger HTT-like malformations, although they have been found to be immature on AVMs and are thought to contributing to the instability of vessels ^60^. In addition, recent reports indicate the potential importance of MCs coverage in treatment of HHT^60,61^. Furthermore, Sugden and colleagues recently also highlighted the importance of haemodynamic forces in this context and demonstrated that ENG function is necessary to mediate blood flow induced EC shape changes which limit vessel diameter and prevent the formation of AV-shunts ^62^. This is in line with our hypothesis, which suggests that blood vessel dilation and AV-shunt formation can be triggered if haemodynamic changes are not counteracted. We propose, that, in *Srf^iMCKO^* mice this is likely caused by the loss of the myogenic response. Our study suggests that vSMC can play a fundamental role in the development of AVMs and might put vSMC in AVM research in a future focus.

## 5. Acknowledgments

We thank Catrin Bitter and Lars Muhl for critically reading the manuscript. Furthermore we thank Caner Bagci for help with the bioinformatic analysis. We are grateful to Anke Biedermann, Siegfried Alberti, and the staff of the UKT animal facility for support with mouse husbandry. We want to thank Christian Feldhaus (light microscopy facility, MPI Tuebingen), Kristin Bieber (FACS facility, UKT) and Gabi Frommer-Kästle (EM facility, UKT) for technical support.

## Supplemental Materials

Expanded Materials & Methods

Online Figures I-VIII

Online Videos I-IV

Online References 63-68

## Non-standard Abbreviations and Acronyms

Actb: Beta actin
AV: arteriovenous
AVMs: arteriovenous malformations
CNS: Central nervous system
DEG: Differential expressed genes
ECs: Endothelial cells
EdU: 5-ethynyl-2’-deoxyuridine
ERG: Electroretinography
FACS: Fluorescence-activated cell sorting
fc: Fold change
FDR: False discovery rate
flex1: Floxed exon 1
GO: Gene ontology
GSEA: Gene set enrichment analysis
ICG: Indocyanine green
iMCKO: Induced mural cell knockout
kbp: Kilo base pair
KEGG: Kyoto Encyclopedia of Genes and Genomes
MCs: Mural cells
MRTF: Myocardin related transcription factor
NG2: Neural/glial antigen 2
NVTs: Neovascular tufts
OIR: Oxygen induced retinopathy
P: Postnatal day
pBPCs: Primary brain pericytes
PCA: Principal Component analysis
PCs: Pericytes
PDGFB: Platelet derived growth factor b
PDGFRB: Platelet derived growth factor receptor beta
RBCs: Red blood cells
SLO: Scanning laser ophthalmoscopy
SMCs: Smooth muscle cells
SMG: Smooth muscle gene
SRF: Serum response factor
TCFs: Ternary complex factors
vSMC: vascular smooth muscle cell
αSMA: Alpha-smooth muscle actin

## 9. Tables

None.

## References

1. Adams RH, Alitalo K. Molecular regulation of angiogenesis and lymphangiogenesis. Nat Rev Mol Cell Biol. 2007;8:464–478.

2. Holm A, Heumann T, Augustin HG. Microvascular Mural Cell Organotypic Heterogeneity and Functional Plasticity. Trends Cell Biol [Internet]. 2018;28:302–316. Available from: http://dx.doi.org/10.1016/j.tcb.2017.12.002

3. Armulik A, Genové G, Mäe M, Nisancioglu MH, Wallgard E, Niaudet C, He L, Norlin J, Lindblom P, Strittmatter K, Johansson BR, Betsholtz C. Pericytes regulate the blood-brain barrier. Nature. 2010;468:557–561.

4. Gaengel K, Genové G, Armulik A, Betsholtz C. Endothelial-mural cell signaling in vascular development and angiogenesis. Arterioscler Thromb Vasc Biol. 2009;29:630–638.

5. Vanlandewijck M, He L, Mäe MA, Andrae J, Ando K, Del Gaudio F, Nahar K, Lebouvier T, Laviña B, Gouveia L, Sun Y, Raschperger E, Räsänen M, Zarb Y, Mochizuki N, Keller A, Lendahl U, Betsholtz C. A molecular atlas of cell types and zonation in the brain vasculature. Nature. 2018;554:475–480.

6. Hammes HP, Lin J, Renner O, Shani M, Lundqvist A, Betsholtz C, Brownlee M, Deutsch U. Pericytes and the pathogenesis of diabetic retinopathy. Diabetes. 2002;51:3107–3112.

7. Eilken HM, Diéguez-Hurtado R, Schmidt I, Nakayama M, Jeong HW, Arf H, Adams S, Ferrara N, Adams RH. Pericytes regulate VEGF-induced endothelial sprouting through VEGFR1. Nat Commun. 2017;8.

8. Park DY, Lee J, Kim J, Kim K, Hong S, Han S, Kubota Y, Augustin HG, Ding L, Kim JW, Kim H, He Y, Adams RH, Koh GY. Plastic roles of pericytes in the blood-retinal barrier. Nat Commun. 2017;8:1–16.

9. Figueiredo AM, Villacampa P, Diéguez-Hurtado R, José Lozano J, Kobialka P, Cortazar AR, Martinez-Romero A, Angulo-Urarte A, Franco CA, Claret M, Aransay AM, Adams RH, Carracedo A, Graupera M. Phosphoinositide 3-kinase-regulated pericyte maturation governs vascular remodeling. Circulation. 2020;142:688–704.

10. Nadeem T, Bogue W, Bigit B, Cuervo H. Deficiency of Notch signaling in pericytes results in arteriovenous malformations. JCI Insight [Internet]. 2020 [cited 2021 May 7];5. Available from: https://pubmed.ncbi.nlm.nih.gov/33148887/

11. Armulik A, Genové G, Betsholtz C. Pericytes: Developmental, Physiological, and Pathological Perspectives, Problems, and Promises. Dev Cell. 2011;21:193–215.

12. Webb RC. Smooth muscle contraction and relaxation. Am J Physiol - Adv Physiol Educ. 2003;27:201–206.

13. Touyz RM, Alves-Lopes R, Rios FJ, Camargo LL, Anagnostopoulou A, Arner A, Montezano AC. Vascular smooth muscle contraction in hypertension. Cardiovasc Res. 2018;114:529–539.

14. Zhou N, Lee JJ, Stoll S, Ma B, Wiener R, Wang C, Costa KD, Qiu H. Inhibition of SRF/myocardin reduces aortic stiffness by targeting vascular smooth muscle cell stiffening in hypertension. Cardiovasc Res. 2017;113:171–182.

15. Wolinsky CD, Waldorf H. Chronic Venous Disease. Med Clin North Am. 2006;93:1333–1346.

16. Miano JM. Serum response factor: Toggling between disparate programs of gene expression. J. Mol. Cell. Cardiol. 2003;35:577–593.

17. Posern G, Treisman R. Actin’ together: serum response factor, its cofactors and the link to signal transduction. Trends Cell Biol. 2006;16:588–596.

18. Olson EN, Nordheim A. Linking actin dynamics and gene transcription to drive cellular motile functions. Nat Rev Mol Cell Biol. 2010;11:353–365.

19. Angstenberger M, Wegener JW, Pichler BJ, Judenhofer MS, Feil S, Alberti S, Feil R, Nordheim A. Severe Intestinal Obstruction on Induced Smooth Muscle-Specific Ablation of the Transcription Factor SRF in Adult Mice. Gastroenterology. 2007;133:1948–1959.

20. Wiebel FF, Rennekampff V, Vintersten K, Nordheim A. Generation of mice carrying conditional knockout alleles for the transcription factor SRF. Genesis. 2002;32:124–126.

21. Chen Q, Zhang H, Liu Y, Adams S, Eilken H, Stehling M, Corada M, Dejana E, Zhou B, Adams RH. Endothelial cells are progenitors of cardiac pericytes and vascular smooth muscle cells. Nat Commun. 2016;7:1–13.

22. Mericskay M, Blanc J, Tritsch E, Moriez R, Aubert P, Neunlist M, Feil R, Li Z. Inducible Mouse Model of Chronic Intestinal Pseudo-Obstruction by Smooth Muscle-Specific Inactivation of the SRF Gene. Gastroenterology. 2007;133:1960–1970.

23. Muzumdar MD, Tasic B, Miyamichi K, Li L, Luo L. A Global Double-Fluorescent Cre Reporter Mouse. Genesis. 2007;605:593–605.

24. Nguyen UTV, Bhuiyan A, Park LAF, Kawasaki R, Wong TY, Wang JJ, Mitchell P, Ramamohanarao K. An automated method for retinal arteriovenous nicking quantification from color fundus images. IEEE Trans Biomed Eng. 2013;60:3194–3203.

25. Franco CA, Jones ML, Bernabeu MO, Geudens I, Mathivet T, Rosa A, Lopes FM, Lima AP, Ragab A, Collins RT, Phng LK, Coveney P V., Gerhardt H. Dynamic Endothelial Cell Rearrangements Drive Developmental Vessel Regression. PLoS Biol. 2015;13:1–19.

26. Ogura S, Kurata K, Hattori Y, Takase H, Ishiguro-Oonuma T, Hwang Y, Ahn S, Park I, Ikeda W, Kusuhara S, Fukushima Y, Nara H, Sakai H, Fujiwara T, Matsushita J, Ema M, Hirashima M, Minami T, Shibuya M, Takakura N, Kim P, Miyata T, Ogura Y, Uemura A. Sustained inflammation after pericyte depletion induces irreversible blood-retina barrier breakdown. JCI Insight. 2017;2:1–22.

27. Lukinavičius G, Reymond L, D’Este E, Masharina A, Göttfert F, Ta H, Gūther A, Fournier M, Rizzo S, Waldmann H, Blaukopf C, Sommer C, Gerlich DW, Arndt HD, Hell SW, Johnsson K. Fluorogenic probes for live-cell imaging of the cytoskeleton. Nat Methods. 2014;11:731–733.

28. Weinl C, Riehle H, Park D, Stritt C, Beck S, Huber G, Wolburg H, Olson EN, Seeliger MW, Adams RH, Nordheim A. Endothelial SRF/MRTF ablation causes vascular disease phenotypes in murine retinae. J Clin Invest. 2013;123:2193–2206.

29. Franco CA, Blanc J, Parlakian A, Blanco R, Aspalter IM, Kazakova N, Diguet N, Mylonas E, Gao-Li J, Vaahtokari A, Penard-Lacronique V, Fruttiger M, Rosewell I, Mericskay M, Gerhardt H, Li Z. SRF selectively controls tip cell invasive behavior in angiogenesis. Dev. 2013;140:2321–2333.

30. Vartiainen MK, Guettler S, Larijani B, Treisman R. Nuclear Actin Regulates Dynamic Subcellular Localization and Activity of the SRF Cofactor MAL. Science (80-). 2007;316:1749–1752.

31. Dubrac A, Künzel SE, Künzel SH, Li J, Chandran RR, Martin K, Greif DM, Adams RH, Eichmann A. NCK-dependent pericyte migration promotes pathological neovascularization in ischemic retinopathy. Nat Commun. 2018;9:1–15.

32. Selvam S, Kumar T, Fruttiger M. Retinal vasculature development in health and disease. Prog Retin Eye Res. 2018;63:1–19.

33. Tual-Chalot S, Oh SP, Arthur HM. Mouse models of hereditary hemorrhagic telangiectasia: Recent advances and future challenges. Front Genet. 2015;5:1–12.

34. Jin Y, Muhl L, Burmakin M, Wang Y, Duchez AC, Betsholtz C, Arthur HM, Jakobsson L. Endoglin prevents vascular malformation by regulating flow-induced cell migration and specification through VEGFR2 signalling. Nat Cell Biol. 2017;19:639–652.

35. Ola R, Künzel SH, Zhang F, Genet G, Chakraborty R, Pibouin-Fragner L, Martin K, Sessa W, Dubrac A, Eichmann A. SMAD4 Prevents Flow Induced Arteriovenous Malformations by Inhibiting Casein Kinase 2. Circulation. 2018;138:2379–2394.

36. Hellström M, Gerhardt H, Kalén M, Li X, Eriksson U, Wolburg H, Betsholtz C. Lack of pericytes leads to endothelial hyperplasia and abnormal vascular morphogenesis. J Cell Biol. 2001;152:543–553.

37. Esnault C, Stewart A, Gualdrini F, East P, Horswell S, Matthews N, Treisman R. Rho-actin signaling to the MRTF coactivators dominates the immediate transcriptional response to serum in fibroblasts. Genes Dev. 2014;28:943–958.

38. Gualdrini F, Esnault C, Horswell S, Stewart A, Matthews N, Treisman R. SRF Co-factors Control the Balance between Cell Proliferation and Contractility. Mol Cell. 2016;64:1048–1061.

39. Wang Z, Wang DZ, Hockemeyer D, McAnally J, Nordheim A, Olson EN. Myocardin and ternary complex factors compete for SRF to control smooth muscle gene expression. Nature. 2004;428:185–189.

40. Tanimoto N, Muehlfriedel RL, Fischer MD, Fahl E, Humphries P, Biel M, Seeliger MW. Vision tests in the mouse: Functional phenotyping with electroretinography. Front Biosci. 2009;14:2730–2737.

41. Geranmayeh MH, Rahbarghazi R, Farhoudi M. Targeting pericytes for neurovascular regeneration. Cell Commun Signal. 2019;17:1–13.

42. Hirunpattarasilp C, Attwell D, Freitas F. The role of pericytes in brain disorders: from the periphery to the brain. J Neurochem. 2019;150:648–665.

43. Brown LS, Foster CG, Courtney JM, King NE, Howells DW, Sutherland BA. Pericytes and neurovascular function in the healthy and diseased brain. Front Cell Neurosci. 2019;13:1–9.

44. Hellström M, Kalén M, Lindahl P, Abramsson A, Betsholtz C. Role of PDGF-B and PDGFR-β in recruitment of vascular smooth muscle cells and pericytes during embryonic blood vessel formation in the mouse. 1999;3055:3047–3055.

45. Östman A, Andersson M, Betsholtz C, Westermark B, Heldin CH. Identification of a cell retention signal in the B-chain of platelet-derived growth factor and in the long splice version of the A-chain. Mol Biol Cell [Internet]. 1991 [cited 2021 May 7];2:503–512. Available from: /pmc/articles/PMC361840/?report=abstract

46. Lindblom P, Gerhardt H, Liebner S, Abramsson A, Enge M, Hellström M, Bäckström G, Fredriksson S, Landegren U, Nyström HC, Bergström G, Dejana E, Östman A, Lindahl P, Betsholtz C. Endothelial PDGF-B retention is required for proper investment of pericytes in the microvessel wall. Genes Dev. 2003;17:1835–1840.

47. Vasudevan HN, Soriano P. SRF Regulates Craniofacial Development through Selective Recruitment of MRTF cofactors by PDGF signaling. Dev Cell. 2014;31:332.

48. Diéguez-Hurtado R, Kato K, Giaimo BD, Nieminen-Kelhä M, Arf H, Ferrante F, Bartkuhn M, Zimmermann T, Bixel MG, Eilken HM, Adams S, Borggrefe T, Vajkoczy P, Adams RH. Loss of the transcription factor RBPJ induces disease-promoting properties in brain pericytes. Nat Commun [Internet]. 2019;10. Available from: http://dx.doi.org/10.1038/s41467-019-10643-w

49. Winkler I, Bitter C, Winkler S, Weichenhan D, Thavamani A, Hengstler JG, Borkham-Kamphorst E, Kohlbacher O, Plass C, Geffers R, Weiskirchen R, Nordheim A. Identification of Pparγ-modulated miRNA hubs that target the fibrotic tumor microenvironment. Proc Natl Acad Sci U S A. 2020;117:454–463.

50. Shi Z, Ren M, Rockey DC. Myocardin and myocardin-related transcription factor-A synergistically mediate actin cytoskeletal-dependent inhibition of liver fibrogenesis. Am J Physiol - Gastrointest Liver Physiol. 2020;318:G504–G517.

51. Lisabeth EM, Kahl D, Gopallawa I, Haynes SE, Misek SA, Campbell PL, Dexheimer TS, Khanna D, Fox DA, Jin X, Martin BR, Larsen SD, Neubig RR. Identification of Pirin as a Molecular Target of the CCG-1423/CCG-203971 Series of Antifibrotic and Antimetastatic Compounds. ACS Pharmacol Transl Sci. 2019;2:92–100.

52. Sun Q, Chen G, Streb JW, Long X, Yang Y, Stoeckert CJ, Miano JM. Defining the mammalian CArGome. Genome Res. 2006;16:197–207.

53. Owens GK, Kumar MS, Wamhoff BR. Molecular regulation of vascular smooth muscle cell differentiation in development and disease [Internet]. Physiol. Rev. 2004 [cited 2021 Jun 20];84:767–801. Available from: www.prv.org

54. Hong KS, Kim K, Hill MA. Regulation of blood flow in small arteries: Mechanosensory events underlying myogenic vasoconstriction [Internet]. J. Exerc. Rehabil. 2020 [cited 2021 Jun 20];16:207–215. Available from: /pmc/articles/PMC7365734/

55. Clifford PS. Local control of blood flow. Am J Physiol - Adv Physiol Educ [Internet]. 2011 [cited 2021 Jun 23];35:5–15. Available from: https://journals.physiology.org/doi/abs/10.1152/advan.00074.2010

56. PC J, M I. Contributions of pressure and flow sensitivity to autoregulation in mesenteric arterioles. Am J Physiol [Internet]. 1976 [cited 2021 Jul 11];231:1686–1698. Available from: https://pubmed.ncbi.nlm.nih.gov/1052803/

57. Kiel JW. The Ocular Circulation. Colloq Ser Integr Syst Physiol From Mol to Funct. 2011;3:1–81.

58. PC J. The myogenic response in the microcirculation and its interaction with other control systems. J Hypertens Suppl [Internet]. 1989 [cited 2021 Jul 11];7. Available from: https://pubmed.ncbi.nlm.nih.gov/2681595/

59. Peacock HM, Caolo V, Jones EAV. Arteriovenous malformations in hereditary haemorrhagic telangiectasia: Looking beyond ALK1-NOTCH interactions. Cardiovasc Res. 2016;109:196–203.

60. Thalgott J, Dos-Santos-Luis D, Lebrin F. Pericytes as targets in hereditary hemorrhagic telangiectasia. Front Genet. 2015;5:1–16.

61. Lebrin F, Srun S, Raymond K, Martin S, Brink S Van Den, Freitas C, Bréant C, Mathivet T, Larrivée B, Thomas J, Arthur HM, Westermann CJJ, Disch F, Mager JJ, Snijder RJ, Eichmann A, Mummery CL. Thalidomide stimulates vessel maturation and reduces epistaxis in individuals with hereditary hemorrhagic telangiectasia. Nat Med. 2010;16:420–429.

62. Sugden WW, Meissner R, Aegerter-Wilmsen T, Tsaryk R, Leonard E V., Bussmann J, Hamm MJ, Herzog W, Jin Y, Jakobsson L, Denz C, Siekmann AF. Endoglin controls blood vessel diameter through endothelial cell shape changes in response to haemodynamic cues. Nat Cell Biol. 2017;19:653–665.

