## Supplemental material for "Mural cell SRF controls pericyte migration, vessel patterning and blood flow"

### Expanded Methods

#### Ethical statement

All animal experiments carried out in this work were performed in compliance with the relevant laws and guide lines of that apply to the University of Tuebingen. Conducted animal experiments were approved by the local animal ethics committee and conducted with the permission by the regional authorities in Tuebingen (Regierungspraesidium Tuebingen), Germany (IM05/18G and IM02/19G).

#### Mouse lines and inducible genetic experiments

*Pdgfrb(BAC)-CreER$^{T2}$* mice [21] were bred with *Srf-flex1* mice [20] carrying a floxed exon 1 (flex1) of the *Srf* gene to obtain *Pdgfrb-CreERT2::Srf-flex1* mice on a C57BL/6 genetic background. For recombination and high resolution microscopy analyses, the *Rosa$^{mTmG}$* reporter gene [23] was crossed into *Pdgfrb-CreERT2::Srf-flex1*. Postnatal gene deletion in newborn mice was induced via inraperitoneally (i.p.) injection of 50 µg tamoxifen (Sigma, #T5648, dissolved in ethanol-peanut oil (#P2144)) into the milkspot (stomach) on postnatal days (P) 1 to P3, respectively. In OIR experiments, animals were injected i.p. with 200 µg of tamoxifen on 3 consecutive days between P12 and P14. To induce Cre-mediated recombination in adult animals (8 weeks old), 500 mg of tamoxifen was injected i.p. on 5 consecutive days. Both males and females were used for genetic experiments and are represented in a 1:1 ratio.

#### Tissue preparation and Immunohistology

Retinas from postnatal mice were isolated for subsequent staining procedures. Whole eye globes were taken and fixed in 4 % paraformaldehyde (PFA, Sigma, #P6148) solved in phosphate buffered saline (PBS) for 2h either on ice (mild fixation for antibody (AB) staining) or at room temperature (RT; strong fixation for IB4 staining). Eye balls were washed in PBS followed by dissection of the retina. Afterwards, the retina was partially cut into four quadrants. Freshly prepared retinas were blocked and permeabilized in blocking solution containing 1 % BSA (Sigma, #A4378) and 0.5 % Triton X-100 (Sigma, #T8787) overnight at 4 °C. Primary AB were dissolved in blocking buffer (supplemented with 2 % donkey serum (Sigma, #S30-M)) and incubated over night at 4 °C. The next day, the primary antibody solution was removed and retina samples were washed 5 times for at least 20 min with PBS. Next, samples were incubated with immunofluorescent species suitable secondary antibodies (see **online table I**) for 2h at RT protected from light. The incubation was followed by 5 washing times as already indicated. Finally, samples were mounted on a microscope slide using Fluoromount G (Southern Biotech, #0100-01) mounting medium.

When an isolectin-B4 (IB4) staining was performed, blocked Retina samples were washed thrice in Pblec buffer (1 µM $MgCl_2$, 1 µM $CaCl_2$ , 0.1 µM $MnCl_2$ and 0.5 % TritonX-100 in PBS) and incubated with IB4 (see **online table I**) overnight in Pblec at 4 °C. When a co-staining was necessary, primary antibodies were additionally added. The next day, IB4 stained tissues were washed 5 times for 20 min with PBS and subsequently incubated in fluorophore coupled Streptavidin (see **online table I**) together with additional secondary antibodies in blocking buffer for 2h at RT. Subsequent washing and mounting procedure were carried out as mentioned above. All mounted retina samples were stored protected from light at 4 °C.

**Transmission electron microscopy**

Eye balls were freshly removed and retinas were dissected without any fixation. After dissection, the retinas were fixed in Karnowsky's fixative (2 % PFA, 2 % glutaraldehyde (Sigma, #G5882) solved in 0.1 M cacodylate buffer). Retinas were then washed in cacodylate buffer and post-fixed in 1 % osmium tetroxide (OsO4, Sigma, #201030) for 1h at RT and afterwards dehydrated in ascending series of ethanol (50, 70, 96, 100 %). In the 70 % step, ethanol was saturated with uranyl acetate (Honeywell Fluka, #73943) for contrast enhancement. Final dehydration was completed in propylene oxide. The specimens were embedded in Araldite adhesive (Serva). Semithin sections (1 µm) and ultrathin sections (50 nm) were generated on a FCR Reichert Ultracut ultramicrotome (Leica). Semithin sections were stained by using toluidine blue. Ultrathin sections were mounted on pioloform-coated copper grids, contrasted with lead citrate and analyzed and documented using an EM10A electron microscope (Carl Zeiss).

**In vivo proliferation assay**

For *in vivo* proliferation of endothelial cells (ECs) thymidine analogue 5-Ethynyl-2´-deoxyuridine (EdU, Invitrogen, #10044) was prepared in DMSO-PBS (10:1, 2 mg/ml) and injected (15 µl/g of body weight) i.p. into P7 and P12 pups 3h before sacrifice. Retinas were prepared and stained as described above. After secondary antibody staining, retinas were stained using the Click-it EdU Alexa Fluor-647 Imaging Kit (Invitrogen, #C10340) according to the manufacturer's instructions.

**Oxygen induced Retinopathy (OIR)**

The OIR disease model was performed as previously described [63] , with slight modifications. In short, neonatal pups and their mothers were kept at normoxic conditions (room air) from birth to P7 to allow normal vascular development. For 5 days from P7 to P12, the mice were transferred to hyperoxic conditions of 75 % $O_2$. Finally, the animals were kept for another 5 days from P12 to P17 under normoxic conditions to induce the pathologic neovascular response. Hyperoxic conditions were realized by transferring the mice to a Biospherix A-Chamber (Biospherix) equipped with a ProOx 110 oxygen controller (Biospherix) connected to technical oxygen (99.99 % $O_2$). To reduce elevated $CO_2$ concentrations in the OIR chamber, 100 g of soda lime was kept at the bottom of the chamber as a quencher. Over the course of the experiment, oxygen concentration in the chamber was controlled twice a day, while health status of the animals was controlled daily. To discharge the animals from the oxygen chamber, the oxygen concentration was decreased stepwise over a time course of 4h (1h of each 60 %, 50 %, 40 %, 30 % $O_2$). Tamoxifen injections to induce Cre-mediated recombination took place from P12 to P14 by 3 consecutive i.p. injections (see above). At P17, pups were sacrificed by exposure to $CO_2$ followed by cervical dislocation. Eye balls were removed and prepared as described above.

#### Fluorescence-activated cell sorting (FACS) of retinal pericytes (PCs)

FACs of retinal PCs was performed as previously described [7]. In short, eye balls from P12 pups were harvested and collected in ice cold MEM (Gibco, #31095-029) supplemented with 25 mM HEPES (Thermofisher, #15630049), penicillin/streptomycin (Gibco, #15140) and 10% fetal calf serum (FCS). Retinas were subsequently dissected and enzymatically digested using the Papain Dissociation Kit (Worthington Biochemical Corporation, #LK003150). After digestion, the single-cell suspension was filtered through a 40 µm cell strainer into a 2 ml reaction tube. Hereafter, the filtered cell suspension was centrifuged for 5 min at 300 x g at 4 °C and the supernatant was carefully removed. The remaining cell pellet was resuspended in 120 FACS buffer (2% FCS and 2mM EDTA pH 8.0). 100 µl of the resuspended cells were mixed with 100 µl of FACS buffer containing fluorescently labeled ABs (Ter119-PB, CD45-PB, CD140a-FITC, CD140b-APC and CD31-PE, final dilutions see **online table I**), followed by incubation for 30 min on ice. The remaining 20 µl of whole retina single-cell suspension was kept as an input control and lysed in 350 µl of RLT buffer plus (Quiagen) supplemented with β-mercaptoethanol (10 µl/ml). After AB incubation, 1 ml of FACS buffer was added to the cell suspension to wash out unbound ABs, followed by centrifugation for 5 min at 300 x g at 4 °C. Finally, the supernatant was carefully removed and the remaining cell pellet was resuspended in 500 µl FACS buffer containing 50 ng/ml DAPI (Sigma, #D9542) to label dead cells. Labeled single cell suspensions were sorted subsequently on a BD FACS Aria lllu sorter (BD Bioscience) using a 100 µm nozzle.

#### Scanning-Laser Ophthalmoscopy (SLO) and optical coherence tomography (OCT)

The *in vivo* analysis of retinal layer structure was performed as published previously [64,65]. In short, mice were anesthetized with ketamine (66.7 mg/kg) and xylazine (11.7 mg/kg) and pupils dilated with tropicamide (Mydriaticum Stulln). OCT imaging was performed together with SLO on a commercially available system (Heidelberg Engineering Spectralis, Eye Explorer version 5.3.3.0). This device features a superluminescent diode at 870 nm as low coherence light source. OCT scans are acquired at a speed of 40.000 scans per second and each two-dimensional B-scan (set to 30° field of view) contains up to 1536 A-scans. We use black on white mode, i.e. high reflectivity is shown as dark and low reflectivity as white. For angiography, indocyanine green dye (ICG) was used as published [6,7] previously in combination with an infrared laser (795 nm; barrier filter 800 nm).

#### Electroretinography (ERG)

ERGs were obtained according to previously described [40.66]. Mice were dark-adapted overnight before the experiments and their pupils were dilated. Anesthesia was induced by subcutaneous injection of ketamine (66.7 mg/kg), xylazine (11.7 mg/kg) and atropine (1 mg/kg). The Espion V6 equipment (Diagnosys LLC) was used for the ERG measurements. Single flash recordings were obtained both under scotopic (dark-adapted) and photopic (light-adapted)) conditions. Light adaptation was accomplished with a background illumination of 30 $cd/m^2$ for 10 min, which stayed on during the course of the photopic recordings. Stimuli were presented with increasing intensities, reaching from 10–3 $cd*s/m^2$ to 30 $cd*s/m^2$, divided into 10 steps of 0.5 and 1 log $cd*s/m^2$. 10 responses were averaged with an inter-stimulus interval (ISI) of 5 s or 17 s (for 1, 3, 10 and 25 $cd*s/m^2$).

**Western Blot**

Western blot was performed to verify SRF ablation in cultured pBMCs at protein levels. pBMC cultures (control and *Srf-KO*) were grown to 70 % confluence on 6-well cell culture plates. Cells were trypsinized, pelleted and resuspended in ice cold RIPA buffer (50 mM Tris-HCl, 150 mM NaCl, 1 % TritonX-100, 0.5 % Sodium deoxycholate, 0.1 % SDS, 1 mM EDTA, pH 8.0), followed by agitation for 30 min at 4 °C. Subsequent to agitation, resuspended cells were centrifuged for 30 min at 10,000 x g at 4 °C. The supernatant containing proteins was then transferred into a fresh tube and the protein concentration was determined by the Bradford assay (Biorad, #5000201). Protein lysates were stored at -20 °C until used. 20 mg of protein were separated by SDS-PAGE and subsequently transferred onto a 0.22 µm polyvinylidene difluoride transfer membrane using a Bio-Rad transfer system for 50 min (2 mA /$cm^2$ of membrane). Following the transfer, membranes were washed in TBST buffer (138 mM NaCl, 20 mM Tris, 0.1% Tween-20 in $H_2O$, pH 7.4) once and subsequently blocked in blocking buffer containing 5% nonfat died milk in TBST for 1h at RT. After blocking, membranes were incubated with the primary AB (see **online table I**) diluted in blocking buffer overnight at 4°C. The next day, the membranes were washed 3 times with TBST and subsequently incubated with corresponding horse radish peroxidase (HRP) coupled secondary AB (see **online table I**) diluted in blocking buffer for 1h at RT. Finally, membranes were washed three times with TBST followed by a washing step with TBS (138 mM NaCl, 20 mM Tris in $H_2O$, pH 7.4) to remove detergents from the membrane. Imaging of membranes was performed by addition of chemiluminescent HRP substrate (Biorad, #1705061). Documentation was performed on a Fusion SL documentation system (Vilber, version V.070).

**Luciferase assay**

NIH/3T3 cells (ATCC CRL-1658) were transfected with SRF-reporter plasmids $(TSM)_2$ and $(TMM)_2$ [28] using the TransIT-LT1 (Mirus Bio LLC, #MIR 2300) transfection reagent. Afterwards, the cells were starved in starvation media (DMEM+0.3 % FCS), or in starvation media containing MRTF inhibitor (CCG-203971, 30 µM; Sigma, #SML1422). The following day, cells were stimulated (excluding control conditions) by addition of Platelet-Derived Growth Factor-BB (PDGFBB; 25 ng/µl; Sigma, #SRP3138) for 7h. Afterwards, media was removed and lysates for luciferase assay were prepared according to the manufacturer's instructions (Promega, #E1910). Luciferase activity was measured using an OPTIMA FluoroSTAR (BMG LABTECH).

**Isolation and culture of primary brain pericytes (pBPCs)**

pBPCs from murine brains Brain mural cells were isolated from 6 to 8 weeks old C57BL/6 mice which harbor the homozygous *Srf-flex1* allele and a heterozygous *Rosa26$^{mTmG}$* allele. The isolation procedure was performed as previously described [67] with minor changes. In short, brains of 3 mice were dissected and placed in ice cold MEM media containing 1%n pen/strep. Using a razor blade, the olfactory bulb, cerebellum and medulla were removed. The remaining cortical tissues were pooled and rigorously minced using a razor blade. Minced brains were washed using MEM and centrifuged for 5 min at 300 x g at RT. The tissue pellet was enzymatically digested using the Papain Dissociation Kit (Worthington Biochemical Corporation, #LK003150) according to the manufactures protocol at 37°C for 70 min. After incubation, the digested tissue was homogenized by slow trituration, passing it 10 times through a 18G needle, followed by passing it 5 times through a 21G needle. The homogenate was then resuspended in 5 ml of 22 % BSA/PBS solution and centrifuged at 1,300 x g for 10 min to separate the cell content in the pellet from the myelin in the upper phase. The supernatant was removed completely by aspiration and the cell pellet was then washed in EBM-2 media (Lonza, #CC-3162) followed by centrifugation at 300 x g for 5 min. After aspiration of the supernatant, the pellet containing single cells and vascular micro vessels was resuspended in 6 ml EBM2-medium and seeded equally in 3 wells of a 6- well plate coated with 0,2 % collagen type I (ColI; Corning, #CB354249). Plated cells were incubated at 37 °C and 5 % $CO_2$ for 20h in a humidified incubator to ensure attachment of viable cells to the plate. After incubation, the plates were carefully washed 3 times with 3 ml PBS. Finally, 3 ml of fresh EMB-2 media were added to each well and the cells were cultured until confluence. Cultures were then trypsinized (Gibco, #25200056), pooled and further cultured in a T25 cell culture flask. The established cell culture contained endothelial cells (ECs), fibroblasts and PCs, was cultured in EBM-2 medium until the second passage. pBPC culture, was established by culture the cells in PC medium (Sciencell, #1201) from the third passage onwards, which especially promotes mural cell growth. From passage 5 onwards, pBPC cultures were judged to have reached high purity [11]. Expression of mural cell markers was determined by qPCR.

**Tat-Cre transduction of pBPCs**

For Tat-Cre transduction, 5 x $10^4$ cells/6-well or 2 x $10^4$ cells/24-well of pBPCs were seeded on 0.2% ColI coated plates and cultured overnight. The next day, cells were washed with PBS followed by addition of OptiMEM (Thermofisher, # 31985062) reduced serum medium. Then, Tat-Cre (Merck, #SCR508) was added to the wells (final concentration 4µM) and incubated with the cells for 20h. The next day, the cells were carefully washed with PBS and recovered in PC-medium for 2 days. After recovery time, cells were directly used for experiments. The recombination efficiency was estimated by counting of eGFP positive cells compared to total cell count. Furthermore *Srf* gene expression was determined by qPCR and western blot analyses.

**Wound closure migration assay**

*Srf-KO* and respective control pBPCs were trypsinized and counted as previously mentioned. 10 x $10^4$ cells were seeded to 4-well plates of a 0.2 % ColI-coated 35 mm µ-Dish (Ibidi, #80466). Cells were cultured overnight in PC-medium and the next day, the culture insert was removed to generate wound-like gaps. The dish was washed with PBS and PC-medium containing 20 ng/ml PDGFBB (Sigma, #SRP3138), 750 nM SiR-Actin (Spirochrome, #SC001) as well as 1 µM verapamil was added. The stained cells were imaged for 20h at 37°C using a LSM800 (Zeiss) microscope at 10x magnification (PApo 10x/0.45) with 1 image cycle/10 min.

**Trans-well migration assay**

*Srf-KO* and respective control pBPCs were trypsinized and counted. Afterwards, the cells were resuspended in starvation media (basal PC-medium supplemented with 0.3% FCS) and 4 x $10^4$ cells were seeded into a Boyden chamber insert with 8 µm pore size placed in a 24-well cell culture plate. Cells were incubated for 2h at 37°C and to let cells adhere. Then, the inserts were transferred to a new well containing 600 µl starvation medium supplemented with 20 ng/ml PDGFBB (Sigma, #SRP3138). Plates were incubated for 6h at 37°C to allow cells to migrate towards the gradient. Migration was stopped by replacement of the medium and subsequent fixation of the membranes using PFA for 10 min at RT. After washing of the membrane with PBS, cells were incubated in DAPI (3 µg/ml) to stain the nucleus of migrated cells. Membranes were subsequently 3 times washed with PBS. To remove not migrated cells, the inner part of the membrane was cleaned by a cotton swab. Finally, the membrane was cut out of the chamber and mounted on a microscope slide. Samples were stored at 4°C until imaged by fluorescence microscopy.

**RNA-Seq analysis**

RNA of FAC sorted cells was isolated using the RNeasy micro plus kit (Quiagen, #74034) according to the manufacturer's instructions and assessed on a 2100 BioAnalyzer (Agilent technologies). RNA-Seq libraries were generated with the Single Cell/Low Input RNA Library Prep Kit (New England biolabs, #E6420L) and sequenced on an Illumina device. The raw sequencing files were obtained in FASTQ format and quality controls were assessed by using FASTQC (https://www. bioinformatics.babraham.ac.uk/projects/fastqc/). Obtained reads were aligned to the mouse genome assembly (GRCm38.p6 Ensembl release 92) using TopHat2 (version 2.1.1), following by generation of gene expression counts by using HTSeq-count (version 0.6.1) with the option -m intersection-nonempty. Final gene expression analysis across the measured samples was performed by using DESeq2 package on protein-coding genes. A principal component analysis (PCA) was used to assess the overall similarity between the samples and performed based on transformed read counts. Differentially expressed genes (DEGs) were selected by using a false discovery rate p-value cut-off 0.5. Gene enrichment analysis (GSEA) of DEGs was performed using the GSEA-tool (version 4.0.3) and a cut-off p-value of <0.05.

**Morphometric analysis of retinal vasculature**

Retinal vasculature microscopy images were analyzed using the ImageJ software tool Fiji (version 1.52p) [68]. In all quantifications, the control group was set as reference and the mutant group was calculated accordingly. All images for quantifications were obtained from two retinas out of one animal and were averaged and expressed as one value (n=1). In general, 8 (but at least 4) comparable fields of view were imaged per animal and averaged. Images for quantification were taken at high magnification (40x) and with a field of view covering 317 µm*317 µm. The number of utilized animals in each group is indicated in the respective figures. Vascularized area was measured as ICAM2$^+$ , CD31$^+$ or IB4$^+$ area divided by total area. Radial outgrowth was measured as distance from the optic nerve to the periphery of the vascular front for each wedge of the retina. Endothelial filopodia at the angiogenic front were manually counted and normalized to a length of 1000 µm. Branching points were manually counted and divided by the vascularized area. Mural cell coverage was determined by manually counting NG2$^+$ or PDGFRB$^+$ cells at the capillary plexus and the angiogenic front, normalized to the vascularized area. Mural cell coverage determined by DES staining was measured by DES$^+$ area divided by the total vascularized area. Empty sleeve area was measured as CollV$^+$ area subtracted by the IB4$^+$ vessel area, leaving only retracted vessels. This retracted vessel area was divided by IB4$^+$ vessel area. SMA signal intensity was measured by SMA intensity on arteries normalized to ICAM2 signal intensity. The background intensity of each assessed channel was subtracted from the respective channel intensity. SMA intensity was assessed by measurement of SMA signal intensity, subtracted by the background intensity. Extravasation of red blood cells was determined by subtraction of the vascularized area from the Ter119$^+$ erythrocyte area to detect extravasated red blood cells. Arteriovenous nicking (crossings of arterioles and venules) was counted manually on whole retina samples. EC density was determined by manually counting ERG$^+$ nuclei to specifically detect ECs and was normalized by dividing by the ICAM2$^+$ vessel area. Levels of proliferating ECs were determined by counting of ERG$^+$/EdU$^+$ double-positive cells, normalized to ERG$^+$ cells. Arterial and venous diameter was measured on three independent positions within a distance of up to 1000 µm to the optic nerve and the individual diameter values were averaged. Vascular smooth muscle cell (vSMC) coverage was measured by manually counting eGFP$^{+}$ cells or measurement of NG2$^{+}$ or DES$^{+}$ area on arteries and veins, normalized to the vascularized area. Avascular area of retinas in OIR experiments was measured by using the "polygon selection" tool and normalized to the vascularized area. NVT area was determined by manual selection of NVT areas using the affinity software (Serif; expressed in pixel values) normalized to the total area. PC and SMA coverage on NVTs was determined by measurement of the NG2$^{+}$ or DES$^{+}$ area normalized to the vascularized area.

#### Statistics

Statistical analyses were performed using the GraphPad Prism 9 software. All data if not otherwise stated are presented as mean ± standard deviation of the mean (s.d.). All experimental *in vivo* data are based on animal numbers (n) derived from at least two different litters. The unpaired two-tailed Students t-test with Welch's correction or the Mann Whitney test were used to determine statistical significance. For comparison of multiple groups a one-way ANOVA and Tukey's post hoc test were used. A *P-value*<0.05 was considered to be statistically significant.

**Online table I. List of antibodies and staining reagents.**

| Antigen/staining reagent | Source, clone/number | Company | Dilution |
|---|---|---|---|
| NG2 | Rabbit, AB5320 | Millipore | 1:100 |
| CD140a-FITC | Rat, APA5, 11-1401-82 | Invitrogen | 1:100 |
| CD140b-APC | Rat, APB5, 17-1402-82 | Invitrogen | 1:25 |
| CD144 | Rat, 11D4.1 ,555289 | BD Biosciences | 1:100 |
| CD31 | Goat,AF3628 | R&D Systems | 1:400 |
| CD31-PE | Rat, 390, 102408 | Biolegend | 1:50 |
| CD45-PB | Rat, 30F-11, 103126 | Biolegend | 1:200 |
| CollV | Rabbit, 2150-1470 | Bio-Rad | 1:200 |
| Desmin | Rabbit, ab15200 | Abcam | 1:200 |
| ERG1 | Rabbit, ab110639 | Abcam | 1:100 |
| GFP-Alexa Fluor 488 | Rabbit, A21311 | Invitrogen | 1:200 |
| ICAM-2 | Rat,553326 | BD Pharmigen | 1:200 |
| Isolectin B4 | Bandeiraea simplicifolia | Sigma-Aldrich | 1:25 |
| PDGFRβ | Goat, AF1042, | R&D Systems | 1:100 |

| | | | |
|---|---|---|---|
| SiR-actin | SC001 | Spirochrome | 1:1000-1:3000 |
| SMA-Cy3 | Mouse,C6198, 1A4 | Sigma-Aldrich | 1:300 |
| SRF | Rabbit, D71A9, 5147S | Cell Signaling Technology | 1:500 |
| Streptavidin Alexa Fluor-400 | S32351 | Invitrogen | 1:100 |
| Streptavidin Alexa Fluor-488 | S11223 | Invitrogen | 1:100 |
| Ter119 | Rat,TER-199,553671 | BD Biosciences | 1:200 |
| Ter119-PB | Rat, TER-119, 116232 | Biolegend | 1:200 |
| α-rab HRP | Goat, 7074 | Cell Signaling Technology | 1:10000 |
| α rat Alexa Fluor-488 | Donkey, A21208 | Invitrogen | 1:500-1:1000 |
| α-goat Alexa Fluor-488 | Donkey, A32814 | Invitrogen | 1:500-1:1000 |
| α-goat Alexa Fluor-647 | Donkey, A21447 | Invitrogen | 1:500-1:1000 |
| α-rab Alexa Fluor-488 | Donkey, A21206 | Invitrogen | 1:500-1:1000 |
| α-rab Alexa Fluor-568 | Donkey, A10042 | Invitrogen | 1:500-1:1000 |
| α-rab Alexa Fluor-647 | Goat, A21244 | Invitrogen | 1:500-1:1000 |
| α-rat Alexa Fluor-647 | Chicken, A21472 | Invitrogen | 1:500-1:1000 |

### Online Figures

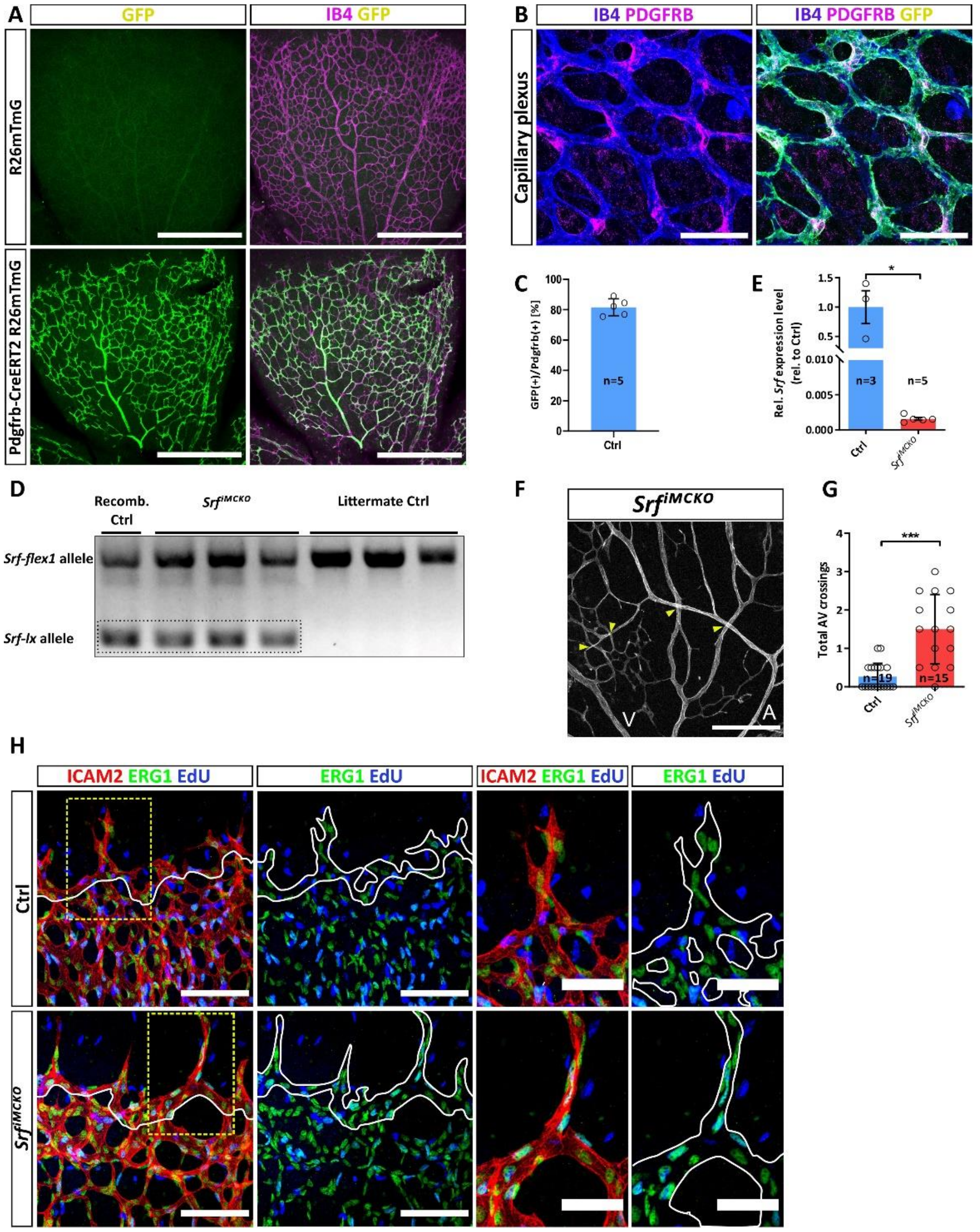


**Online Figure I| Verification of *Srf* deletion in retinal MCs and extended morphometric analyses**. **(A)** Confocal images of tamoxifen induced *Pdgfrb-Cre*$^{ERT2}$::*Srf-flex1* heterozygous mice that carry the *Rosa26*$^{mTmG}$ reporter . eGFP expression (green) in mural cells confirms the specificity of *Pdgfrb-Cre*$^{ERT2}$. Endothelial cells were co-stained with IB4. Scale bar, 500 µm. **(B)** High resolution confocal images of retinas stained for endothelial cells (IB4, blue), mural cells (PDGFRB, magenta) and recombinant cells (eGFP, green), highlighting specific recombination in mural cells. Scale bar, 50 µm. **(C)** Quantification of the recombination rate of MCs. **(D)** Generic PCR of whole retina lysates from *Srf*$^{iMCKO}$ mice showing the efficient recombination of the *flex1*-allele upon tamoxifen administration, resulting in a *Srf-lx* (exon 1 deletion) band. In littermate control mice, only the unrecombined *flex*1-allele becomes amplified **(E)** qPCR quantification of *Srf* expression levels of FACS sorted retinal MCs. (**F-G**) Confocal image and respective quantification of arteriovenous (AV)-nicking events (yellow arrowheads) in *Srf*$^{iMCKO}$ retinas ("A" indicates artery and "V" vein). Scale bar, 200 µm. **(H)** Extended figure of figure 1(J) showing confocal images of proliferating cells (EdU, blue), endothelial cell apical surface (ICAM2, red) and endothelial nuclei (ERG1, green). White lines indicate the border of the leading front (first column) or the vessel shape (second and forth column). Images on the right panels show magnification of the yellow dashed squares shown on the left panel. Scale bar, 100 µm (left) and 50 µm (right). Error bars indicate s.d. of the mean. Statistical comparison by unpaired t-test with Welch's correction. Number of analyzed animals or independent repetitions (n) are indicated. ns= not significant, *p≤0.05, **p≤0.01, ***p≤0.001.

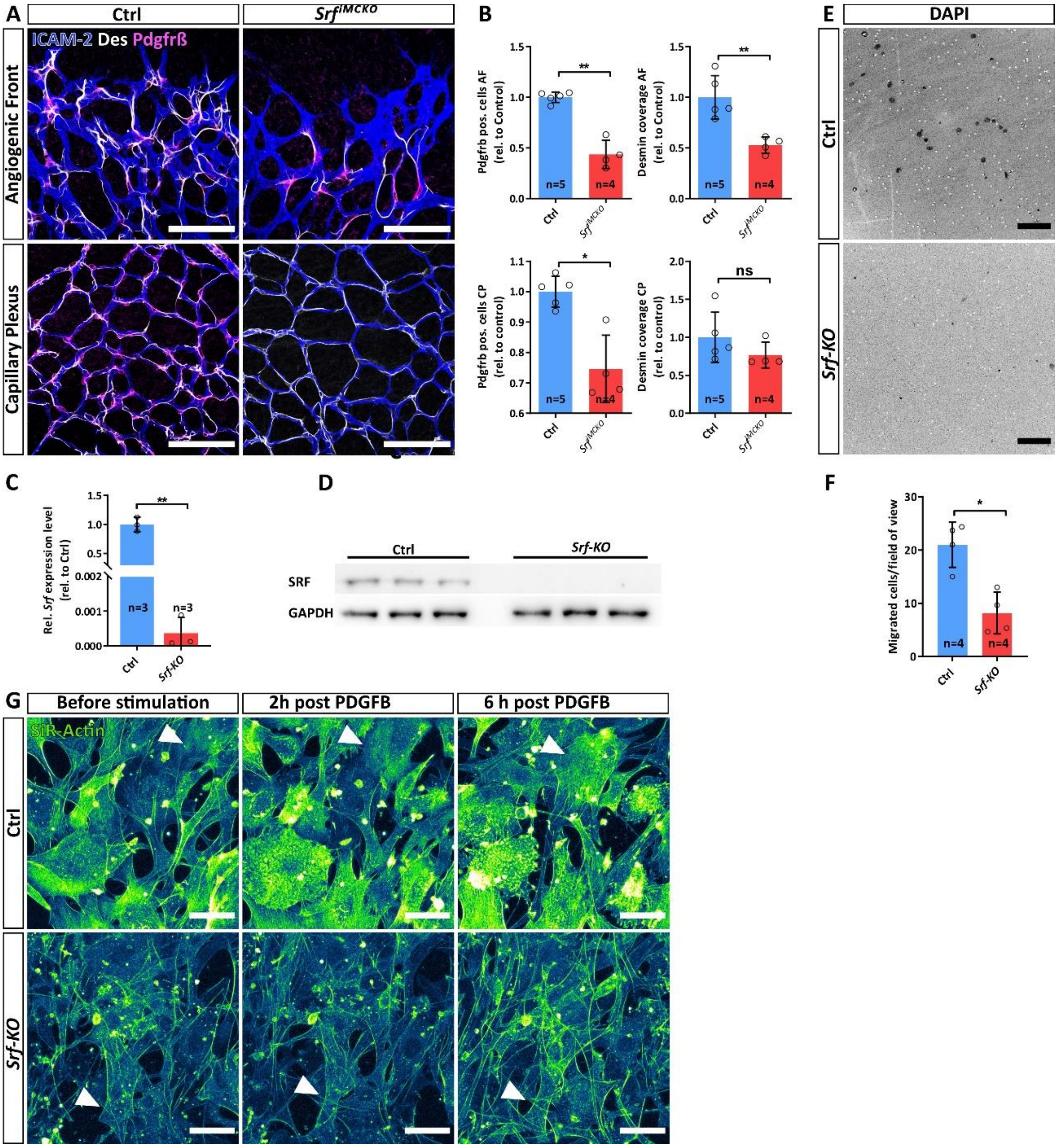


**Online Figure II| SRF-deficient pericytes (PCs) show migration defects**. **(A)** Confocal images showing PC coverage at the capillary plexus (CP) and the angiogenic front (AF). ECs were stained with ICAM2 (blue) and PCs with desmin (white) and PDGFRB (magenta). Scale bar, 100 µm. **(B)** Respective quantifications of the PC coverage. **(C, D)** Verification of genetic Srf delition in Tat-Cre treated primary Srf-flex1 pBPCs via qPCR and westernblot analyses, respectively. **(E)** Confocal images of a trans-well assay to determine polarized migration towards a PDGFB gradient trough a membrane with defined pore size. DAPI-positive dots (black) indicate migrated cells which passed the membrane. Scale bar, 100 µm **(F)** Respective quantification showing the average of migrated cells per field of view. **(G)** Confocal images of a time-laps experiment showing stimulation of Ctrl and Srf-KO pBPCs with PDGFB. Note the accumulation of F-actin in Ctrl cells (white arrowheads, Ctrl row), whereas the Srf-KO cells show no detectable accumulation of F-actin (white arrowheads, Srf-KO row). Scale 50 µm. Error bars show s.d. of the mean. Statistical comparison by unpaired t-test with Welch's correction. Number of analyzed animals/repetitions (n) is indicated. ns = not significant, *p≤0.05, **p≤0.01, ***p≤0.001.

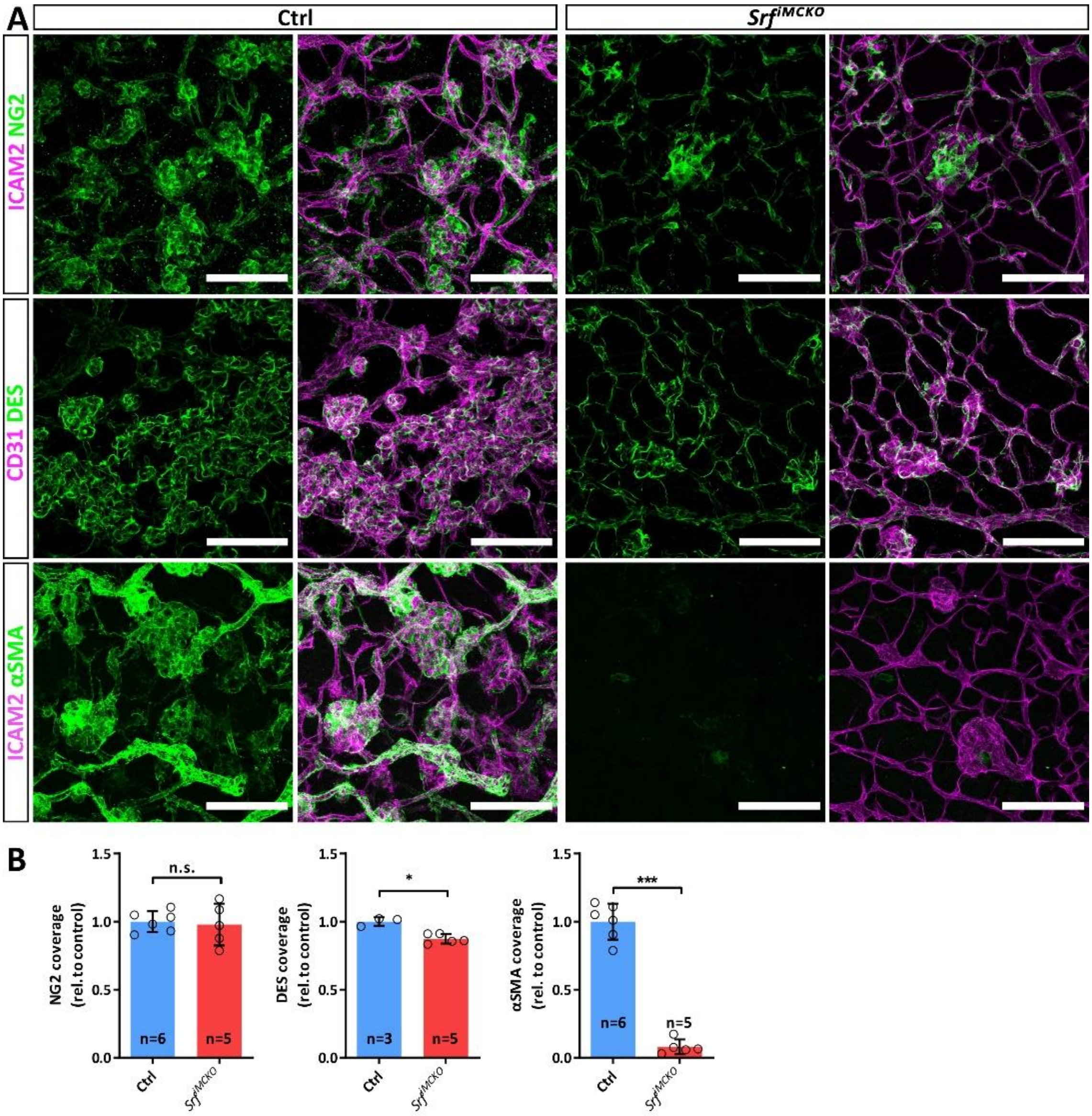


**Online Figure III| SRF is a driving force of pathologic pericyte (PC) activation. (A)** Confocal images of control and *Srf*$^{iMCKO}$ OIR retinas showing NVTs. Staining of NG2 (green, first row) or DES (green, second row) indicate PCs covering ICAM2 (magenta, first row) or CD31 (magenta, second row) stained vessels. Note the loss of αSMA signal (green, third row) as well as the reduction of amount and size of neovascular tufts (NVTs) in *Srf*$^{iMCKO}$ OIR retinas. Scale bar, 100 µm. **(B)** Quantification of PC coverage of NVTs, using NG2 and DES as markers and of pathologically PC activation determined by αSMA coverage. Error bars indicate s.d. of the mean. Statistical comparison by unpaired t-test with Welch's correction. Number of analyzed animals (n) is indicated. ns = not significant, *p≤0.05, **p≤0.01, ***p≤0.001.

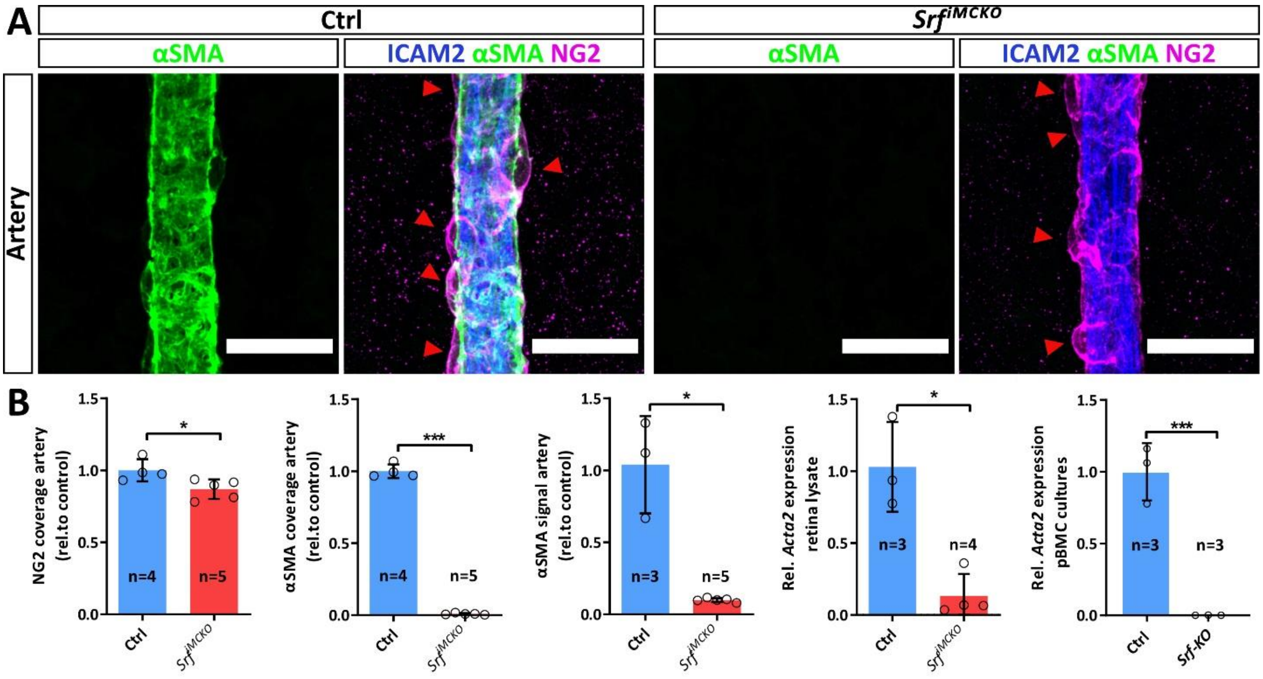


**Online Figure IV| *$Srf^{iMCKO}$* vascular smooth muscle cells (vSMCs) fail to express the contractile muscle protein αSMA. (A)** Confocal high-resolution images of P6 arteries, showing endothelial cells (ECs) (ICAM2, blue), vSMC marker αSMA (green) and mural cell (MC) marker NG2 (magenta). Red arrowheads indicate colocalization of αSMA and NG2. Note the abolishment of αSMA in *$Srf^{iMCKO}$*. Scale bars, 25 µm. **(B)** Quantification of aSMC coverage determined by NG2 and αSMA coverage as well as by the measured intensity of αSMA staining. Relative transcript level of *Acta2* (encoding αSMA) determined by qPCR analysis of whole retina and pBPC culture lysates. Error bars indicate s.d. of the mean. Statistical comparison by unpaired t-test with Welch's correction. Number of analyzed animals or independent repetitions (n) is indicated. ns = not significant, *p≤0.05, **p≤0.01, ***p≤0.001.

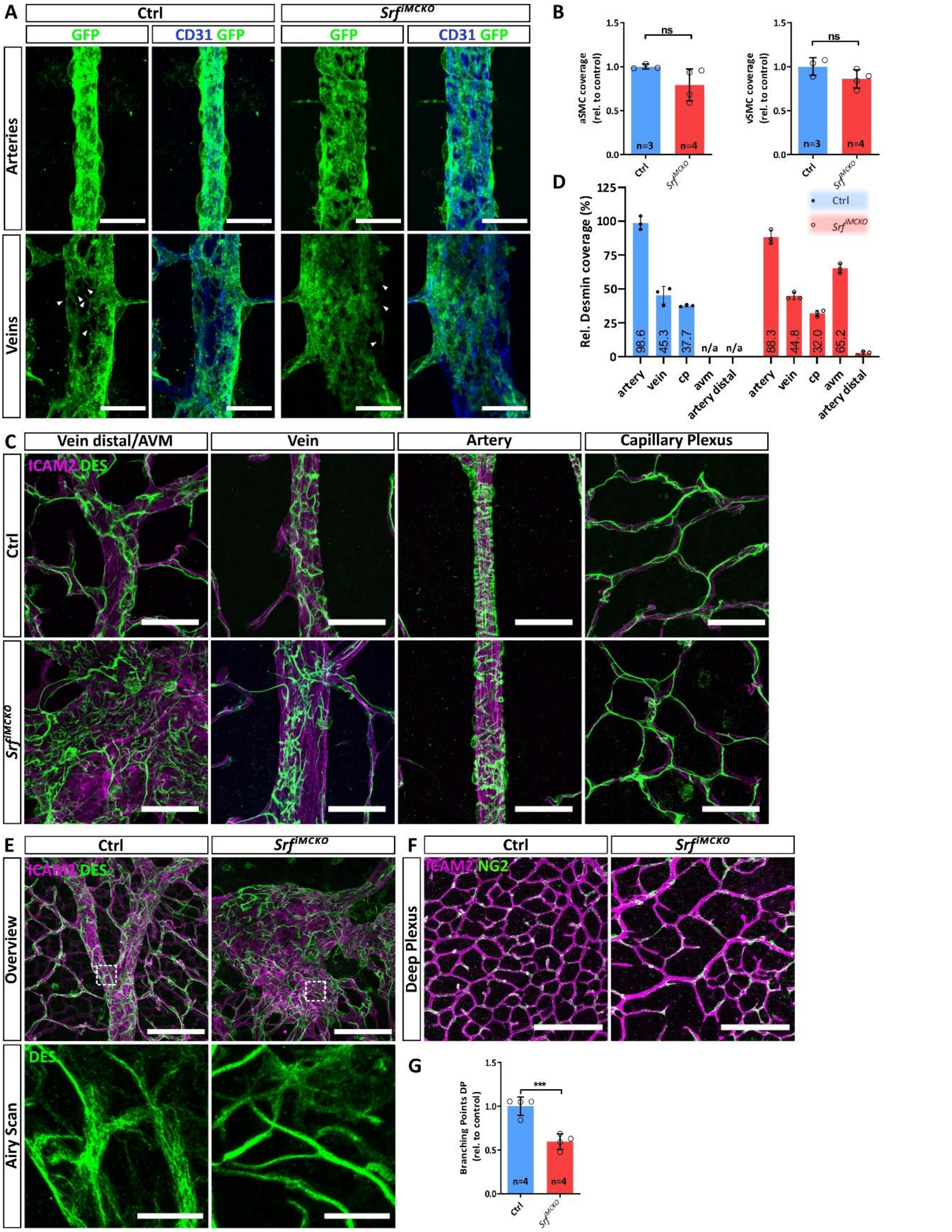


**Online Figure V| Altered morphology of vascular smooth muscle cells (vSMCs) on arteries and veins of $Srf^{iMCKO}$ retinas at P12.** **(A)** Confocal images of retinal arteries and veins labeled with the $Rosa26^{mTmG}$ reporter for mural cells (MCs) (GFP, green) and co-stained for CD31 (endothelial cells, blue). Confocal high-resolution images of retinal arteries and veins. Scale bar, 25 µm **(B)** Quantification of arterial and venous SMC coverage, determined by cell counts per vascular area. **(C)** Confocal images showing representative MC (desmin, green) coverage on vessels (ICAM2, magenta). Scale bar, 50 µm. **(D)** Quantification of MC coverage (determined by desmin area/ICAM2 area). **(E)** Super resolution image (airy scan) of cytoskeletal desmin structure in control and $Srf^{iMCKO}$ vSMCs. **(F)** Vasculature of the deep plexus in P12 control and $Srf^{iMCKO}$ retinas. **(G)** Quantification of branching points of the deep plexus. Error bars indicate s.d. of the mean. Statistical comparison by unpaired t-test with Welch’s correction. Number of analyzed animals (n) is indicated. ns = not significant, *p≤0.05, **p≤0.01, ***p≤0.001.

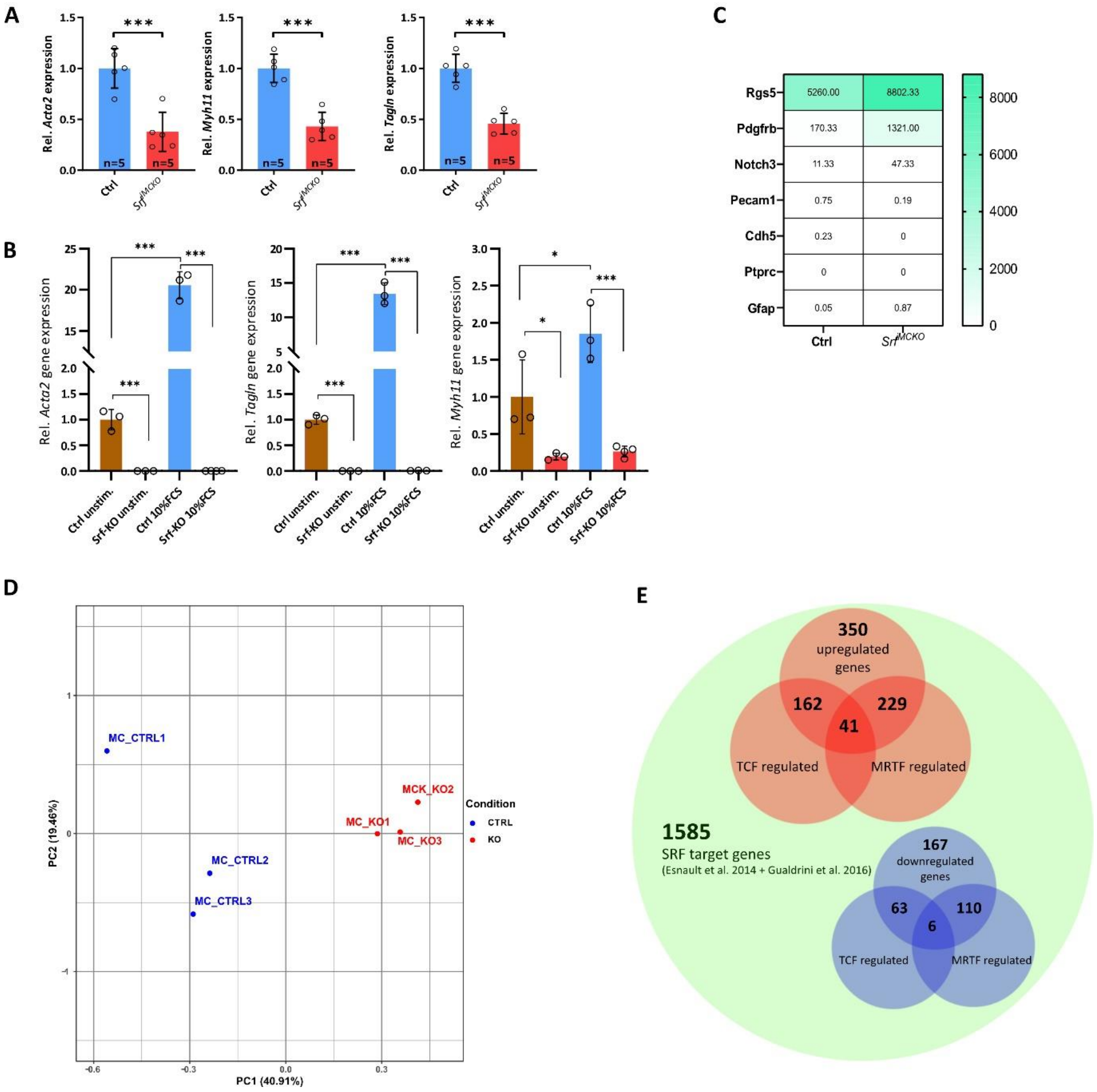


**Online Figure VI| SRF deficient vascular smooth muscle cells (vSMCs) lose expression of contractile genes. (A)** Gene expression of the vSMC markers *Acta2*, *Myh11* and *Tagln* in whole retina lysates. Error bars indicate s.d. of the mean. Statistical comparison by unpaired t-test with Welch’s correction. Number of analyzed animals (n) is indicated. ns = not significant, *p≤0.05, **p≤0.01, ***p≤0.001. **(B)** Transcription levels of vSMC genes *Acta2*, *Tagln* and *Myh11* measured by qPCR. Control and *Srf-KO* pBPCs were maintained in 2 % serum or stimulated with 10 % serum for 24h. Error bars indicate s.d. of the mean. Statistical comparison by one-way ANOVA (Tukey’s multiple comparison test). Number of independent repetitions (n) = 3. ns = not significant, *p≤0.05, **p≤0.01, ***p≤0.001. **(C)** Heatmap of selected EC, MC, leucocyte and astrocyte marker expression across of isolated MCs measured by RNA-Seq. **(D)** Principal component analysis (PCA) plot of RNA-Seq data sets of control and *Srf*$^{iMCKO}$ MCs. **(E)** Venn diagram summarizing upregulated and downregulated SRF target genes detected by RNA-Seq.

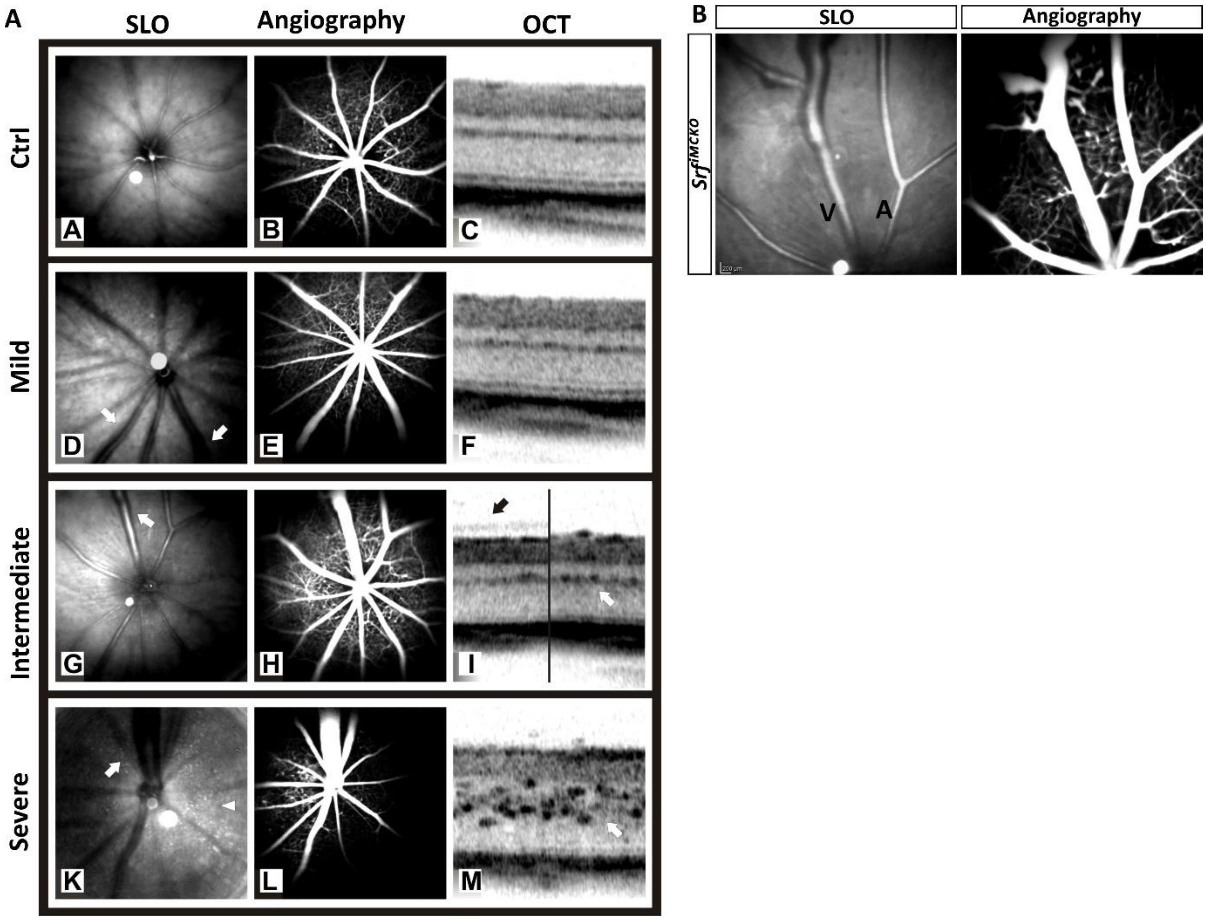


**Online Figure VII| In vivo Scanning laser ophthalmoscopy (SLO)/angiography/ optical coherence tomography (OCT) and ERG measurement. (A)** Extended version of Figure 7 (A) showing SLO, angiography and OCT images of 4 weeks old Ctrl and $Srf^{iMCKO}$ live imaged retinas. **(B)** Higher magnification of $Srf^{iMCKO}$ malformed vessels.

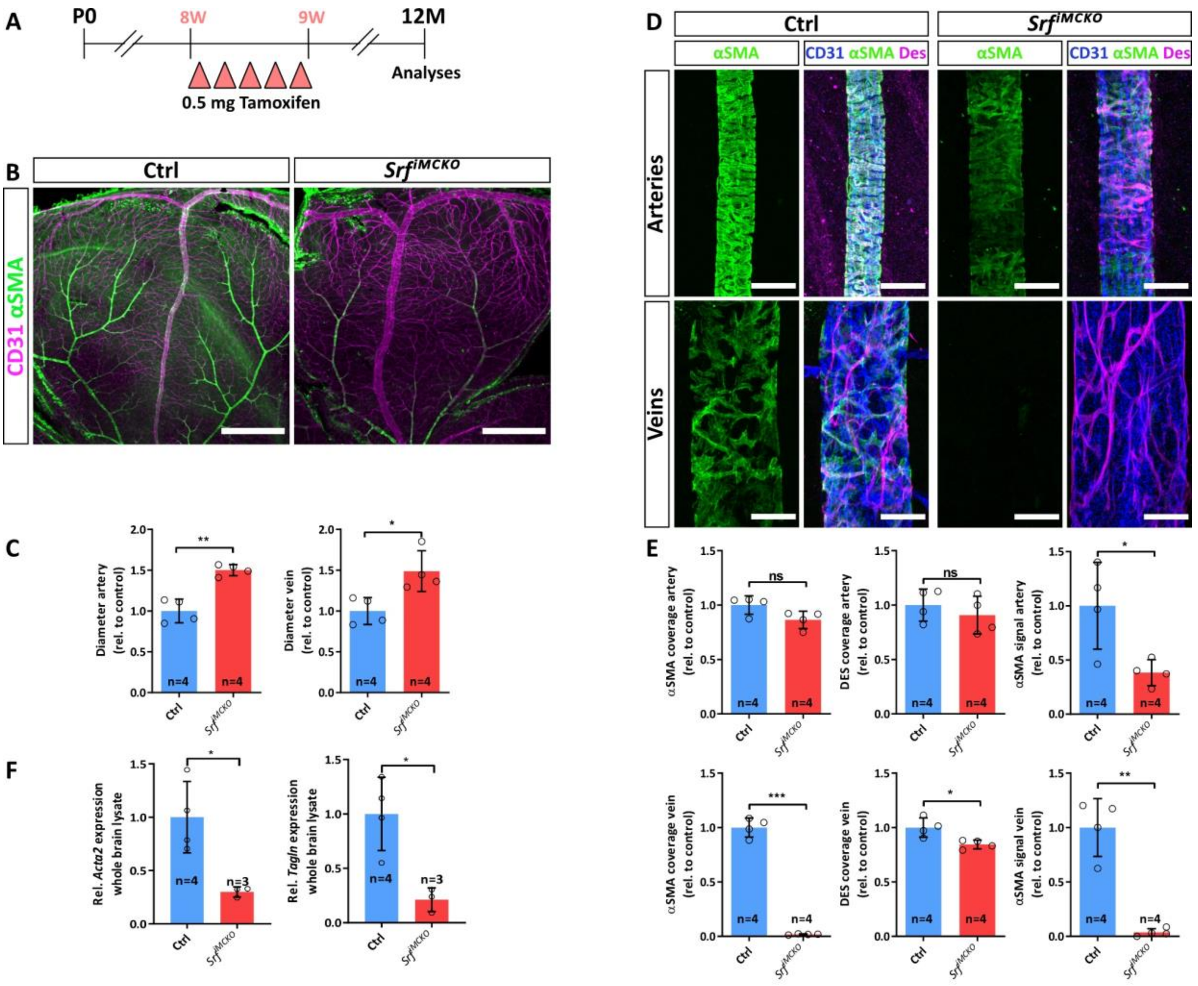


**Online Figure VIII| SRF is necessary for sustained contractile properties of vascular smooth muscle cells (vSMCs). (A)** Illustration of the experimental approach to delete the *Srf* gene in mural cells of adult mice (8-weeks old) via tamoxifen administration at indicated timepoints. Mice were analyzed at the age of 12 months. **(B)** Overview images of the retinal vasculature of control and *Srf*$^{iMCKO}$mice stained for CD31 and αSMA. Note the enlarged vessel diameter of arteries and veins as well as the reduced signal intensity of αSMA staining in *Srf*$^{iMCKO}$ retinas. **(C)** Quantification of diameters of arteries and veins (upper graphs) as well as the quantification of the expression of vSMC marker genes *Acta2* and *Tagln* determined in whole brain lysates. **(D)** Confocal high-resolution images of arteries and veins stained for CD31, αSMA and DES. Note the strong reduction of αSMA signal in *Srf*$^{iMCKO}$. DES positive structures confirm the presence of vSMCs. **(E)** Quantification of αSMA and DES coverage as well as the αSMA signal intensity on arteries and veins. Error bars indicate s.d. of the mean. Statistical comparison by unpaired t-test with Welch's correction. Number of analyzed animals (n) is indicated. ns = not significant, *p≤0.05, **p≤0.01, ***p≤0.001

**Online Videos**

**Online Video I |Representative migration assay of *Srf-KO* and control pBPC cultures stained by SiR-Actin.** Cells were stimulated with PDGFB to stimulate migration. Note the difference in migration speed.

**Online Video II |Representative PDGFB stimulation of *Srf-KO* and control pBPC cultures stained by SiR-Actin.** Cells were stimulated with PDGFB to observe cytoskeletal modulation. Note the absence of reaction of *Srf-KO* pBPCs.

**Online Video III |Representative PDGFB stimulation of 3T3 cells expressing MRTFA-GFP.** 3T3 cells were stimulated with PDGFB to observe MRTFA-GFP nuclear translocation.

**Online Video IV |Representative SLO *in vivo* imaging of $Srf^{iMCKO}$ and respective control retinas**. Note the strong vessel motion of dilated vessels indicating a loss of vascular tone in $Srf^{iMCKO}$ animals.

**Online References**


63. Connor KM, Krah NM, Dennison RJ, Aderman CM, Chen J, Guerin KI, Sapieha P, Stahl A, Willett KL, Smith LEH. Quantification of oxygen-induced retinopathy in the mouse: A model of vessel loss, vessel regrowth and pathological angiogenesis. *Nat Protoc*. 2009;4:1565–1573.

64. Huber G, Beck SC, Grimm C, Sahaboglu-Tekgoz A, Paquet-Durand F, Wenzel A, Humphries P, Michael Redmond T, Seeliger MW, Dominik Fischer M. Spectral domain optical coherence tomography in mouse models of retinal degeneration. *Investig Ophthalmol Vis Sci*. 2009;50:5888–5895.

65. Fischer MD, Huber G, Beck SC, Tanimoto N, Muehlfriedel R, Fahl E, Grimm C, Wenzel A, Remé CE, van de Pavert SA, Wijnholds J, Pacal M, Bremner R, Seeliger MW. Noninvasive, in vivo assessment of mouse retinal structure using optical coherence tomography. *PLoS One* [Internet]. 2009 [cited 2020 Aug 20];4. Available from: https://pubmed.ncbi.nlm.nih.gov/19838301/

66. Tanimoto N, Sothilingam V, Seeliger MW. Functional Phenotyping of Mouse Models with ERG [Internet]. In: Methods in molecular biology (Clifton, N.J.). Methods Mol Biol; 2012 [cited 2020 Aug 20]. p. 69–78.Available from: https://pubmed.ncbi.nlm.nih.gov/23150360/

67. Tigges U, Welser-Alves J V., Boroujerdi A, Milner R. A novel and simple method for culturing pericytes from mouse brain. *Microvasc Res*. 2012;84:74–80.

68. Schindelin J, Arganda-Carreras I, Frise E, Kaynig V, Longair M, Pietzsch T, Preibisch S, Rueden C, Saalfeld S, Schmid B, Tinevez JY, White DJ, Hartenstein V, Eliceiri K, Tomancak P, Cardona A. Fiji: An open-source platform for biological-image analysis. *Nat Methods*. 2012;9:676–682.
